# Systematic prioritization of candidate genes in disease loci identifies *TRAFD1* as a master regulator of IFNγ signalling in celiac disease

**DOI:** 10.1101/2020.03.04.973487

**Authors:** Adriaan van der Graaf, Maria Zorro, Annique Claringbould, Urmo Vosa, Raul Aguirre-Gamboa, Chan Li, Joram Mooiweer, Isis Ricano-Ponce, Zuzanna Borek, Frits Koning, Yvonne Kooy-Winkelaar, Ludvig Sollid, Shuo-Wang Qiao, BIOS consortium, Vinod Kumar, Yang Li, Lude Franke, Sebo Withoff, Cisca Wijmenga, Serena Sanna, Iris Jonkers

## Abstract

**Background:** Celiac disease (CeD) is a complex T cell–mediated enteropathy induced by gluten. Although genome-wide association studies have identified numerous genomic regions associated with CeD, it is difficult to accurately pinpoint which genes in these loci are most likely to cause CeD.

**Results:** We used four different *in silico* approaches – Mendelian Randomization inverse variance weighting, COLOC, LD overlap and DEPICT – to integrate information gathered from a large transcriptomics dataset. This identified 118 prioritized genes across 50 CeD-associated regions. Co-expression and pathway analysis of these genes indicated an association with adaptive and innate cytokine signalling and T cell activation pathways. 51 of these genes are targets of known drug compounds and likely druggable genes, suggesting that our methods can be used to pinpoint potential therapeutic targets. In addition, we detected 172 gene-combinations that were affected by our CeD-prioritized genes in *trans*. Notably, 41 of these *trans*-mediated genes appear to be under control of one master regulator, *TRAFD1*, and were found to be involved in IFNγ signalling and MHC I antigen processing/presentation. Finally, we performed *in vitro* experiments that validated the role of *TRAFD1* as an immune regulator acting in *trans*.

**Conclusions:** Our strategy has confirmed the role of adaptive immunity in CeD and revealed a genetic link between CeD and the IFNγ signalling and MHC I antigen processing pathways, both major players of immune activation and CeD pathogenesis.

## Introduction

Celiac disease (CeD) is an auto-immune disease in which patients experience severe intestinal inflammation upon ingestion of gluten peptides. CeD has a large genetic component, with heritability estimated to be approximately 75%^1^. The largest CeD-impacting locus is the HLA region, which contributes approximately 40% of CeD heritability^2^. While the individual impacts of CeD-associated genes outside the HLA region are smaller, they jointly account for an additional 20% of heritability. Previous genome-wide association studies (GWAS) have identified 42 non-HLA genomic loci associated with CeD^3–6^, but the biological mechanisms underlying the association at each locus and the genes involved in disease susceptibility are largely unknown. Yet, identification of these non-HLA genetic components and an understanding of the molecular perturbations associated with them are necessary to understand CeD pathophysiology.

Understanding the biological mechanisms of non-HLA CeD loci is difficult: only three of these loci point to single nucleotide polymorphisms (SNPs) located in protein-coding regions^3^. The other CeD-risk loci cannot be explained by missense mutations, making it necessary to look at other biological mechanisms such as gene expression to explain their role in CeD pathogenicity. Several studies have been performed to integrate expression quantitative trait loci (eQTLs) with CeD GWAS associations^4,7,8^, and several candidate genes, including *UBASH3A*, *CD274*, *SH2B3* and *STAT4*^9^, have been pinpointed, implicating T cell receptor, NFκB and interferon signalling pathways as biological pathways associated with CeD pathology. Unfortunately, these eQTL studies had limited sample sizes, which reduced their power to identify *cis*- and (especially) *trans*-eQTLs. Furthermore, previous attempts to integrate eQTLs have mostly annotated genomic loci based on catalogued eQTLs without formally testing the causality of the genes in the onset or exacerbation of CeD^8,10,11^.

Gene expression and GWAS data can also be integrated using methodologies that identify shared mechanisms between diseases. These methods can be roughly divided into three classes: variant colocalization methods, causal inference methods and co-expression methods. Colocalization methods consider the GWAS and eQTL summary statistics at a locus jointly and probabilistically test if the two signals are likely to be generated by the same causal variant^12^. Causal inference methods test if there is a causal relationship between expression changes and the disease, using genetic associations to remove any confounders^13,14^. Finally, co-expression methods do not use eQTL information, but rather test if there is significant co-expression between the genes that surround the GWAS locus^15^. Unfortunately, there is no current “gold standard” method for finding the causal gene behind a GWAS hit, as all the methods discussed here are subject to their respective assumptions, drawbacks and caveats. However, it is worthwhile to use all these methods in parallel to find the most relevant causal genes for CeD.

Here, we systematically applied all four methods to the latest meta-analysis results for CeD^5^ and coupled them with eQTL results from the Biobank Integrative Omics Study (BIOS) cohort^16^, one of the largest cohorts for which there is genotype and RNA-seq expression data of peripheral blood mononuclear cells (schematic overview **Fig. 1A-B**). We focused on 58 GWAS loci that showed significant association with CeD at p < 5×10^−6^. Our approach prioritized 118 genes in 50 loci and identified one gene, *TRAFD1*, as a master regulator of *trans*-effects. We then experimentally validated the role of *TRAFD1*-mediated genes using RNA-seq in a disease-relevant cell type. Our study yields novel insights into the genetics of CeD and is proof-of-concept for a systematic approach that can be applied to other complex diseases.

**Fig. 1.**
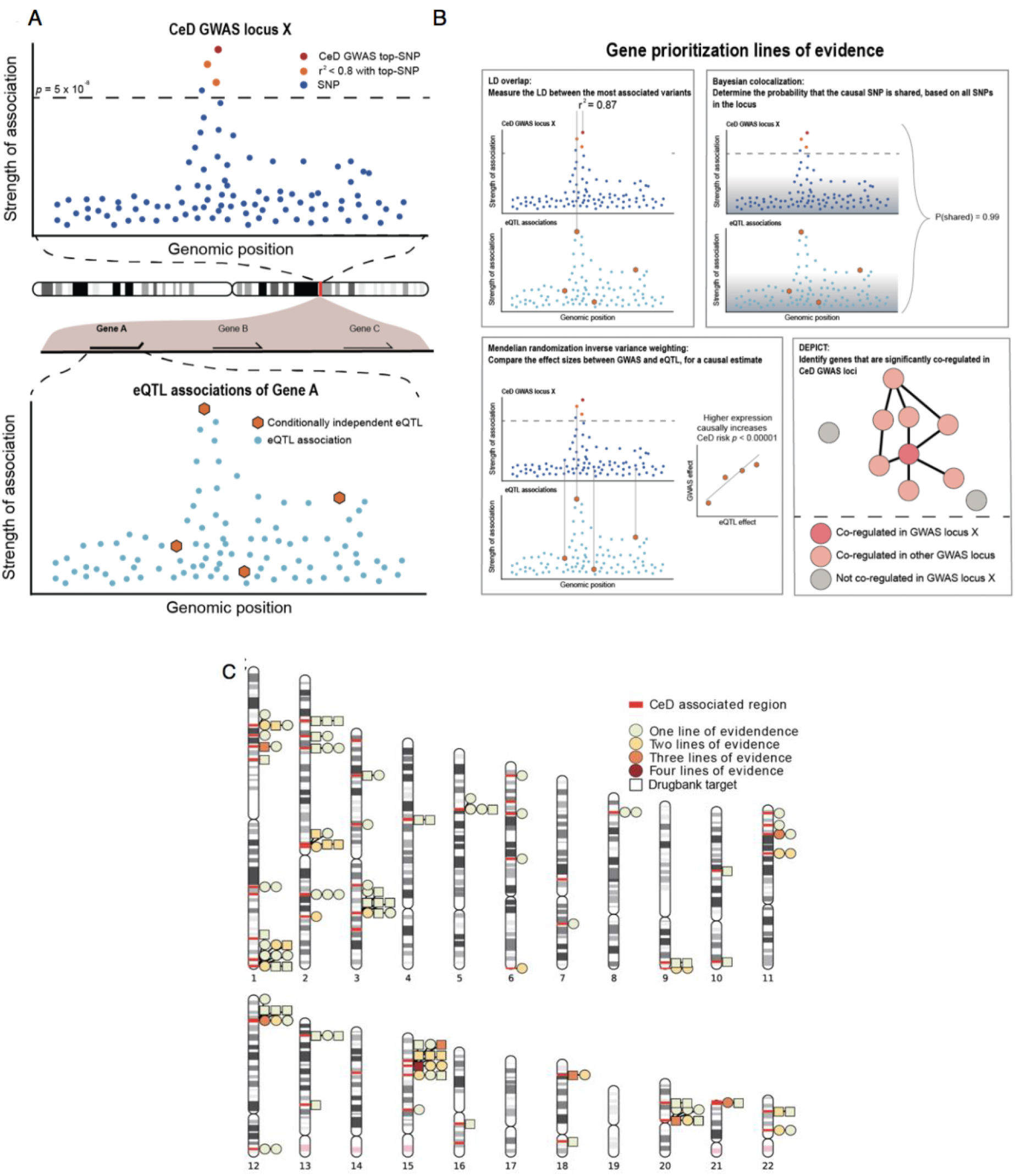
*Cis*-eQTL prioritized candidate genes in CeD loci. (**A**) A CeD GWAS association curve at a hypothetical GWAS locus × and the eQTL association at a potential candidate gene A. In both association plots, each dot represents a SNP plotted against the genomic position (X axis) and the strength of association (Y axis). In the GWAS association curve, the top SNP is marked in red, while other SNPs above the significance threshold (dashed line) are coloured according to their LD with the top SNP. In the eQTL association curve, independent eQTLs are marked in red. (**B**) A conceptual depiction of the four statistical methods applied to link a disease locus to an eQTL locus. (**C**) A chromosome ideogram depicting the location of each prioritized gene identified in a CeD-associated GWAS locus. Loci are marked with red bars. Genes depicted by a square are the target of an approved drug or a drug in development. All other genes are depicted by a circle. Each circle or square is coloured according to the lines of evidences supporting its causal role.

## Methods

### Genotypes for eQTL analysis

We used the BIOS cohort^16^ to map eQTLs in 3,746 individuals of European ancestry. The BIOS cohort is a collection of six cohorts: the Cohort on Diabetes and Atherosclerosis Maastricht^17^, the Leiden Longevity Study^18^, Lifelines DEEP^19^, the Netherlands Twin Registry^20^, the Prospective ALS Study Netherlands^21^ and the Rotterdam Study^22^. As described in Vosa et al.^23^, each cohort was genotyped separately using different arrays. Genotypes were subsequently imputed to the Haplotype Reference Consortium panel (HRC v1.0) on the Michigan imputation server^24^.

We considered only biallelic SNPs with a minor allele frequency (MAF) > 0.01, a Hardy-Weinberg test *p* value > 10^−6^ and an imputation quality RSQR > 0.8. To remove related individuals, a genetic relationship matrix (GRM) was created using plink 1.9^25^ (command –make-grm-bin) on linkage disequilibrium (LD)–pruned genotypes (option: “-- indep 50 5 2”). Pairs of individuals with a GRM value > 0.1 were considered related, and one individual was removed from each of these pairs. Population outliers were identified using a principal component analysis on the GRM, and we removed individuals who were more than 3 standard deviations from the means of principal component 1 or 2.

### Expression quantification

We used the same procedure for RNA gene expression control and processing as described in Zhernakova et al.^16^ In brief, RNA was extracted from whole blood and paired-end sequenced using the Illumina HiSeq 2000 machine. Read alignment of RNA-seq reads was done using STAR (v2.3.0)^26^ using a reference genome with masked variants with MAF < 0.01 in the Genome of the Netherlands^27^. Aligned reads were quantified using HTSeq^28^. Samples were removed if they had fewer than 80% aligned reads, fewer than 85% exon-mapping reads, or if they had a median 3’ bias larger than 70% or smaller than 45%. Unobserved expression confounders were removed following the procedure of Zhernakova et al.^16^, correcting the expression matrix for the first 25 principal components as well as 3’ bias, 5’bias, GC content, intron base pair percentage and sex.

### eQTL analysis

After genotype and RNA-seq quality controls (QCs), 3,503 individuals, 19,960 transcripts and 7,838,327 autosomal SNPs remained for analyses. We performed genome-wide eQTL mapping for the transcripts using plink 1.9^25^ with the --assoc command. We defined *cis*-eQTL variants as those located within ±1.5Mb of the transcript and *trans*-eQTLs as variants located outside these boundaries. To select variants that could explain the *cis*-eQTL signal of a gene, we used GCTA-COJO software^29^ v1.26. For this analysis, we required selected variants to reach a *p*-value threshold of 5 × 10^−6^ and included the BIOS cohort genotypes as LD reference. This identified 707 genes with at least one eQTL reaching this threshold, 357 of which had more than one conditionally independent eQTL variant.

### CeD summary statistics associated regions and candidate genes

We used summary statistics from a CeD GWAS meta-analysis of 12,948 cases and 14,826 controls that analysed 127,855 variants identified using the Immunochip array^5^. SNP positions were lifted over to human genome build 37 using the UCSC liftover tool. We first identified lead associated variants in the CeD meta-analysis by performing *p*-value clumping: we used plink 1.9^25^ to select variants at a *p*-value threshold of 5 × 10^−6^ and pruned variants in LD with these selected variants using standard plink settings (R^2^ > 0.5, utilizing 1000 Genomes European sample LD patterns)^25,30^. We removed variants in an extended HLA region (chromosome 6, 25Mb to 37Mb) due to the complex long range LD structure in this region and because we aim to understand the function of the non-HLA genetic component of CeD. We looked for candidate genes around the clumped variants as follows. First, we defined regions around every clumped variant by padding the clumped SNP position by 1Mb on both sides. We then joined all overlapping CeD-associated regions together and looked for gene transcripts that partly or fully overlapped with the associated regions. This approach identified 58 CeD-associated regions and 1,235 candidate genes that are potentially causal for CeD. Of note, the CeD-association windows were set to be smaller than the eQTL window so that eQTL associations would fully overlap the associated CeD GWAS peak even when a gene is on the edge of the CeD-associated region.

### Gene prioritization using Mendelian Randomization–Inverse Variance Weighting(MR-IVW), COLOC, LD overlap and DEPICT

We prioritized CeD-associated genes using three eQTL-based methods – MR-IVW^31^, COLOC^12^ and LD overlap – and one co-regulation-based method, DEPICT^15^. For the MR-IVW method, we used the independent variants identified by GCTA-COJO as instrumental variables^13,32^ to test causal relationships between changes in gene expression and CeD. MR-IVW was only performed when there were three or more independent eQTLs available (164 genes). A gene was significant for the MR-IVW test if the causal estimates passed a Bonferroni threshold *p*-value of 3.1 × 10^−4^. Heterogeneity of causal estimates was accounted for and corrected for using Weighted Median MR analysis and Cochran’s Q test^33^. For the COLOC method, we used the ‘coloc’ R package and considered a gene significant for the COLOC analysis if the posterior probability of shared variants (H4) was larger than 0.9. For the LD overlap method, a gene was considered significant if there was high LD (r^2^ > 0.8) between the top independent eQTL and the top CeD variant in the region. Finally, we applied DEPICT^15^ to the clumped CeD GWAS variants described in ‘CeD summary statistics associated regions and candidate genes’. Genes identified by the DEPICT analysis were considered significant if a False Discovery Rate (FDR) < 0.05 was found with DEPICT’s own permutation measure.

We scored each gene in the CeD-associated loci by considering each of the four prioritization methods. A gene was prioritized as ‘potentially causal’ in CeD pathology when one of the four methods was significant (one line of evidence). If multiple lines of evidence were significant, the gene was prioritized more highly than when only a single line of evidence was available.

To explore how the prioritized genes affect CeD risk, we gave each gene an effect direction based on the effect direction of the top variants in the eQTL and the CeD GWAS using the following algorithm:

1. If there was a concordant effect that was significant in the top variants of both the eQTLs and the GWAS, the direction of the concordant effect was chosen.
2. If there was a concordant effect, but no significance of the SNP in one of the datasets, we could not be sure of an effect direction, and a question mark was chosen. The only exception to this was if the MR-IVW was significant, when we chose the direction of the MR-IVW effect.
3. If there was a discordant effect between the top SNPs, and both were significant in both datasets, a question mark was chosen. The only exception to this was when the IVW was significant, when the IVW effect was chosen.
4. If there was a discordant effect and there was significance in only one of the GWAS from the eQTL top SNP, the eQTL direction was chosen.
5. If there was a discordant effect and there was significance in only one of the eQTL from the GWAS top SNP, the GWAS direction was chosen.
6. If there was otherwise a discordant effect, a question mark was chosen.

Each gene is given a mark: positive (‘+’), negative (‘−’) or unknown (‘?’). ‘+’ indicates that increased expression increases CeD risk. ‘−’ indicates that increased expression decreases CeD risk. ‘?’ indicates that it is unknown how the expression affects CeD risk.

### Co-regulation clustering

The genes that have been prioritized may have some shared function in CeD pathology. To identify possible shared pathways, we performed co-regulation clustering analysis based on 1,588 normalized expression co-regulation principal components identified from RNA-seq information across multiple human tissues by Deelen et al^34^. We performed pairwise Pearson correlation of our prioritized genes with these 1,588 principal components and derived a correlation Z score for each prioritized gene pair. We then performed hierarchical clustering of this Z score matrix using Ward distances and identified 4 clusters from the resulting dendrogram.

### *Trans* eQTL and mediation analysis

238 autosomal genes that were not located in, but were associated with, a significant *trans*-eQTL variant (*p* < 5×10^−8^) in the CeD-associated regions were identified and used as potential targets for mediation by our associated genes in the CeD-associated loci (86 potential *cis* mediating genes). We first selected *trans*-eQTL genes that were co-expressed (Pearson r > 0.1, 197 gene combinations) with prioritized genes, then performed mediation analysis by running the *trans*-eQTL association again using the expression of the *cis*-eQTL gene as a covariate. We defined a *trans*-mediated gene if, after mediation analysis, the change (increase or decrease) in the effect size of the top *trans*-eQTL variant was significant according to the statistical test described in Freedman and Schatzkin^35^. For this analysis, we used a Bonferroni-adjusted *p*-value of 3.0 × 10^−4^.

### Cell type proportion and *SH2B3* expression mediation analysis

To assess if the *cis*-eQTL effect of *TRAFD1* was not a proxy for cell-type composition, we performed mediation analyses in a fashion similar to the *trans* mediation analysis above using cell proportions measured in a subset of individuals in the BIOS cohort. To ensure that there was no residual effect of *SH2B3*-expression on the mediating effect of *TRAFD1*, we corrected the original *TRAFD1* expression levels for the expression levels of *SH2B3*, leaving *TRAFD1* expression independent of *SH2B3*, and reran the mediation analysis.

#### Literature review

We performed a REACTOME pathway^36^ analysis to determine the potential function of the prioritized genes. This was complemented with a literature search (research and review papers) in Pubmed. For the coding and non-coding genes for which no studies were found, Genecards (www.genecards.org) and Gene Network v2.0 datasets (www.genenetwork.nl)^34^ were used, respectively. Information regarding the potential druggability of the prioritized genes was obtained from DrugBank^37^, the pharmacogenetics database^38^ and a previous study that catalogued druggable genes^39^.

#### THP-1 culture

The cell line THP-1 (Sigma Aldrich, ECACC 88081201) was cultured in RPMI 1640 with L-glutamine and 25mM HEPES (Gibco, catalogue 52400-025), and supplemented with 10% fetal bovine serum (Gibco, catalogue 10270) and 1% penicillin/ streptomycin (Lonza, catalog DE17602E). The cells were passed twice per week at a density lower than 0.5 × 10^6^ cells/ml in a humidified incubator at 5% CO_2_, 37°C.

#### siRNA treatment

THP-1 cells were plated at 0.6 × 10^6^ cells/ml and transfected with 25 nM siRNA using Lipofectamine RNAimax transfection reagent (Invitrogen, catalogue 13788), according to the manufacturer’s protocol. Cells were treated with an siRNA to target *TRAFD1* (Qiagen catalogue 1027416, sequence CCCAGCCGACCCATTAACAAT) (Knockdown (KD)), and cells treated with transfection mix without siRNA (Wild type (WT)) or non-targeting control siRNA (scrambled (SCR)) (Qiagen catalog SI03650318, sequence undisclosed by company) were included as controls. All the treatments were performed in triplicate. 72 hours after transfection, a small aliquot of cells was stained for Trypan Blue exclusion to determine cell viability and proliferation. The cells were stimulated with either LPS (10 ng/ml) from *E. coli* (Sigma catalogue 026:B6) or media alone (unstimulated) for 4h. Subsequently, the cells were centrifuged, and the cell pellets suspended in lysis buffer and stored at −80°C until used for RNA and protein isolation.

#### qPCR

The total RNA from THP-1 cells was extracted with the mirVana™ miRNA isolation kit (AMBION, catalogue AM1561) and subsequently converted to cDNA using the RevertAid H Minus First Strand cDNA Synthesis Kit (Thermo scientific, catalogue K1631). qPCR was done using the Syber green mix (Bio-Rad, catalogue 172-5124) and run in a QuantStudio 7 Flex Real-Time system (Applied Biosystems, catalogue 448598). Primer sequences to determine KD levels of *TRAFD1* were 5’ GCTGTTAAAGAAGCATGAGGAGAC and 3’ TTGCCACATAGTTCCGTCCG. *GAPDH* was used as endogenous qPCR control with primers 5’ ATGGGGAAGGTGAAGGTCG and 3’ GGGGTCATTGATGGCAACAATA. Relative expression values of *TRAFD1* were normalized to the endogenous control *GAPDH* and calculated using the ΔΔCT method, then given as a percentage relative to SCR expression levels.

#### Western blot (WB)

Cell pellets from THP-1 cells were suspended on ice-cold lysis buffer (PBS containing 2% SDS and complete protease inhibitor cocktail (Roche, catalog 11697498001)). Protein concentration of cell extracts was determined using the BCA protein kit (Pierce, catalog 23225). Proteins were separated on 10% SDS-polyacrylamide electrophoresis gel and transferred to a nitrocellulose membrane. After 1 hour of blocking with 5% fat-free milk in Tris-Tween-Buffer-Saline, the membranes were probed for 1 hour at room temperature with mouse mono-clonal TRAFD1 antibody 1:1000 (Invitrogen, catalog 8E6E7) or mouse monoclonal anti-actin antibody 1:5000 (MP Biomedicals, catalog 08691001), followed by incubation with goat anti-mouse horseradish peroxidase–conjugated secondary antibodies 1:10000 (Jackson Immuno Research, catalog 115-035-003). After three 10-minute washes, the bands were detected by Lumi light WB substrate (Roche, catalogue 12015200001) in a Chemidoc MP imaging system (Bio-Rad) and quantified using Image Lab™ software (Bio-Rad). The band intensity of TRAFD1 was normalized to actin, and the TRAFD1 SCR control level was set as 100%.

#### Statistical analysis for *in vitro* experiments in THP-1 cells

The statistical analyses of proliferation, qPCR and WB were performed using Prism 5 software (GraphPad Software, Inc.). Results are presented as mean ± SEM from a representative experiment. Statistical differences were evaluated using a one-tailed *t*-test.

#### RNA sequencing (RNA-seq) in THP-1 cells

RNA from THP-1 cells was extracted with the mirVana™ miRNA isolation kit (AMBION, catalog AM1561). Prior to library preparation, extracted RNA was analysed on the Experion Stdsend RNA analysis kit (Bio-Rad, catalog 7007105). 1 μg of total RNA was used as input for library preparation using the quant seq 3’ kit (Lexogen, catalog 015.96), according to the manufacturer’s protocol. Each RNA library was sequenced on the Nextseq500 (Illumina). Low quality reads, adaptors and poly-A tail reads were removed from FASTQ files. The QC-ed FASTQ files were then aligned to the human_g1k_v37 Ensembl Release 75 reference genome using HISAT default settings^40^, and sorted using SAMtools^41^. Gene-level quantification was performed by the featurecounts function of the RSubread R package v1.6.2^42^. A modified Ensembl version 75 gtf file mapping only to the last 5’ 500 bps per gene was used as gene-annotation database to prevent counting of reads mapping to intra-genic A-repeats. Gene-level differential expression analysis between conditions was performed using the DESeq2 R package^43^ after removing genes with zero counts. Differentially expressed genes (DEGs) were defined as genes presenting an absolute log2 fold change (|log2 FC|) >1 and an FDR ≤ 0.01 across treatment (WT vs. SCR or KD unstimulated cells). To identify the genes responding to LPS stimulation, the DEGs between unstimulated samples and their respective stimulated sample were determined. Venn diagrams were used to depict the relationship between these genes. REACTOME pathway analyses were performed to identify biological processes and pathways enriched in different sets of DEGs using the enrichr API. Enrichments were considered significant if they were below a 0.05 FDR-threshold defined by the enrichr API^36^.

#### Gene set permutation analysis

It can be difficult to determine if a set of genes is ‘on average’ more or less differentially expressed due to co-expression between the genes within the set. To mitigate this, we performed a permutation test that considers the median absolute T statistic calculated by DESeq2^43^ in the WT-SCR experiment as a null observation and compared this null observation with the SCR-KD experimental comparison. This allowed us to compare the expected differential expression of a set of genes, based on the WT-SCR comparison, with the observed differential expression of the same set of genes in the SCR-KD comparison, while still incorporating the co-expression structure of the data. To do this, we randomly selected a same-sized set of genes 1,000,000 times in each relevant experiment (WT-SCR or SCR-KD), and determined the observed median absolute T statistic. We calculated a ratio of how often the permuted value is higher than the observed value. For example, the observations can be that 1% of permuted gene sets are more differentially expressed in the WT-SCR experiment, while only 0.01% of permuted genes sets are more differentially expressed in the SCR-KD experiment. Finally, we divide these values by one another, (percentage SCR-KD)/(percentage WT-SCR), to calculate a fold increase in differential expression. In the example given above, this indicates that the KD is 100 times (0.01/1 = 100) more differentially expressed than expected.

#### Available RNA-seq datasets

Four available RNA-seq datasets were included to study the pattern of expression of prioritized genes. A brief description of each dataset is provided below. (GEO submission in process).

##### Whole biopsy samples

Duodenal biopsies were obtained from 11 individuals (n=6 CeD patients and n=5 controls) who underwent upper gastrointestinal endoscopy (previously described)^44^. All individuals gave informed consent. To identify DEGs between patients and controls, a filter of |log2 FC|>1 and FDR ≤ 0.05 was applied using the DESeq2 R package.

##### Intra epithelial cytotoxic lymphocytes (IE-CTLS)

CD8^+^ TCRαβ IE-CTLs cell lines were isolated from intestinal biopsies and expanded for 12 days, as described previously^45^. Cells were left unstimulated (controls) or treated for 3 hours with IFNβ (300 ng/ml, Pbl Assay science, cat 11410-2), IL-15 (20 ng/ml, Biolegend, cat 570304) or IL-21 (3 ng/ml, Biolegend, cat 571204) (n=8 samples per condition, as previously reported)^44^. Differential expression analysis between unstimulated cells and cytokine-treated IE-CTLs was performed using the R package DESeq2. DEGs were defined as genes presenting a |log2 FC| > 1 and an FDR ≤ 0.05 between untreated controls and cytokine-treated samples.

##### Gluten specific (gs) CD4+ T cells

gsCD4^+^ T cell lines were generated from intestinal biopsies and expanded for 2 weeks, as reported previously^46^. Cells were stimulated for 3 hours with 2.5 μg/ml of anti-CD3 (Biolegend, catalog 317315) and anti-CD28 (Biolegend, catalog 302923) antibodies. Untreated cells were included as control. N=22 samples per condition. DEGs were extracted with the DESeq2 package using the cut-off of |log2fc|>1 and FDR ≤ 0.05 between unstimulated samples and controls.

##### Caco-2 cells

After 2 weeks of expansion in Transwells, the cells were treated with 60 ng/ml of IFNγ (PeproTech) for 3 hours. Untreated cells were included as controls. RNA samples were extracted and further processed for RNA-seq (as described previously^44^). DEGs between control and stimulated cells were extracted with the DESeq2 R package using a cut-off of |log2FC|>1 and FDR ≤ 0.05.

## Results

### Gene prioritization identifies 118 likely causal CeD genes

To identify genes that most likely play a role in CeD (prioritized genes), we combined a recent genome-wide association meta analysis^5^ with (1) eQTLs derived from whole-blood transcriptomes of 3,503 Dutch individuals^16^ and (2) a co-regulation matrix derived from expression data in multiple different tissues and 77,000 gene expression samples^15^. We selected 1,258 genes that were within 1Mb of the 58 CeD-associated non-HLA variant regions (*p* < 5×10^−6^) (see Methods), and prioritized the genes that are the most likely causally related to CeD using four different gene prioritization methods: MR-IVW^13^, COLOC^12^, LD overlap and DEPICT^15^ (**Fig. 1A-B**) (**Supplementary Table 1**).

The first method we applied, MR-IVW, is a two-sample Mendelian Randomization approach called inverse variance weighting (see **Methods**). Our MR-IVW used summary statistics from two datasets: the eQTL and CeD GWAS. First, independent eQTLs at a locus were identified (see **Methods**), then the effect sizes of the eQTL and the GWAS were combined to identify gene expression changes that are causal (or protective) for CeD^13,14^ (see **Methods**). We only applied this method to a subset of 162 genes for which at least three independent *cis*-eQTL variants (at *p* < 5×10^−6^) were identified (see **Methods**)^32^. We accounted for heterogeneity using the Q test and weighted median method and found that the effect sizes were very similar before and after correction (**Supplementary Table 2**).

The second method, COLOC, is a variant colocalization test in which we used eQTL and CeD summary statistics for all the SNPs in a locus and Bayesian probability to infer whether the eQTL and the CeD-association signals are likely to originate from the same causal variant^12^.

The third method, LD overlap, is a more classical annotation-type approach that prioritizes a gene if the top eQTL is in strong LD (r^2^ > 0.8) with the variant most significantly associated with CeD in a locus. This and the COLOC method were applied to 707 genes for which at least one significant eQTL variant was found.

Finally, we used DEPICT^15^, a gene-prioritization method based on co-regulation in expression datasets across multiple different tissues. DEPICT identifies enrichment for co-regulated genes from genes in a GWAS locus. In contrast to the other methods, DEPICT assessed the potential role of all 1,258 genes independently of the presence of an eQTL.

In total, 118 out of the 1,258 assessed genes were prioritized by at least one of the four methods. Of these 118 genes, 28 had two lines of evidence, 7 genes (*CD226*, *NCF2*, *TRAFD1*, *HM13*, *COLCA1*, *CTSH, UBASH3A*) had three lines of evidence, and one gene (*CSK*) was supported by all four methods (**Supplementary Table 1**) (**Fig. 1C**). Overall, we identified potentially causal genes in 50 out of 58 CeD-associated regions.

The four different gene prioritization methods complement each other in different ways. DEPICT prioritized the most genes: 66 in total, 38 of them uniquely prioritized (38/66, 58% unique). One reason for this is that DEPICT is based on co-expression, not genetic background. Indeed, 16 genes prioritized by DEPICT do not have a significant eQTL associated with them. Overall, the most concordance was found between COLOC and LD overlap (30% and 26% unique genes, respectively) as these methods are the most similar, while MR-IVW uniquely prioritized a relatively large proportion of genes (9/20, 45% unique). Thus, each method helps prioritize genes with multiple lines of evidence, but also adds a unique set of genes based on the assumptions of the method.

To see if any of these genes could lead to therapeutic intervention in CeD, we searched for the CeD-associated genes in DrugBank and assessed their druggability potential following Finan et al.^39^ (**Supplementary Table 3**). 26 of the 118 prioritized genes encode proteins that are targeted by an approved drug or a drug in development according to drugbank (**Fig. 1C**) (**Supplementary Table 3**). For example, drugs such as Natalizumab and Basiliximab that target the proteins encoded by *ITGA4* and *IL21R*, respectively, are currently approved or under study for the treatment of immune-mediated diseases including rheumatoid arthritis^47^, Crohn’s disease^48^ and multiple sclerosis^49^ or as an immune-suppressor to avoid kidney transplant rejection. An additional 25 genes encode proteins that are similar to proteins targeted by already approved drugs following Finan et al.^39^ (**Supplementary Table 3**).

### Co-expression patterns of *cis*-eQTL-prioritized loci reveal four functional clusters

The biological function for the 118 prioritized genes and their role in CeD pathology is not fully understood. We sought to infer biological function using a guilt-by-association co-regulation approach to identify clusters of shared molecular function (see Methods). We identified co-regulated genes by correlating our prioritized gene list in 1,588 principal components that were identified from the co-expression of 31,499 RNA-seq samples across multiple tissues^34^ (Fig. 2A). We then performed REACTOME 2016 gene set enrichment^36^ analysis to investigate the biological processes enriched in each cluster (**Supplementary Table 4**) (**Supplementary Table 5**).

**Fig. 2.**
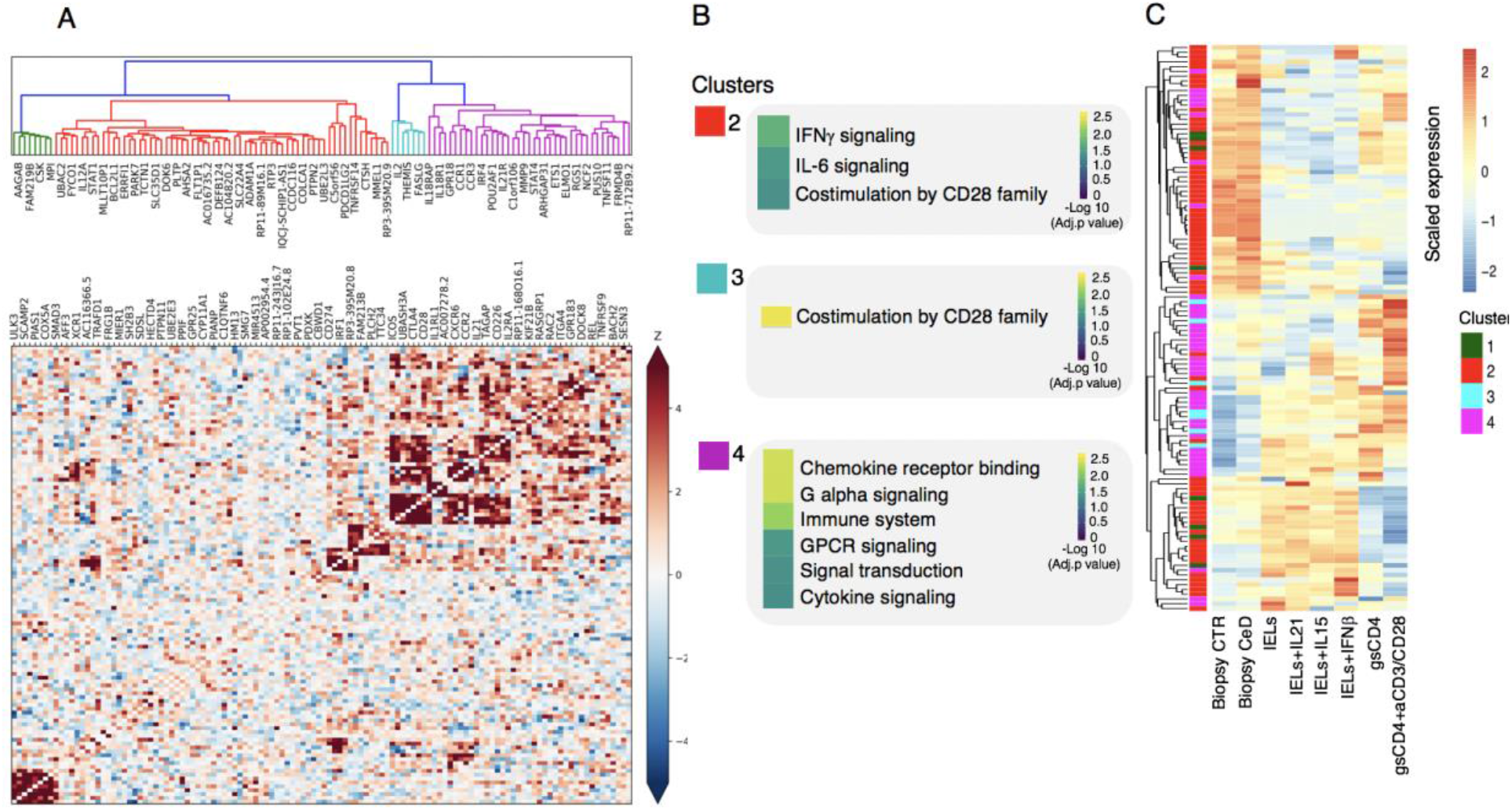
Co-expression pattern of *cis*-eQTL prioritized genes reveals four functional clusters. (**A**) Heatmap showing the Spearman correlations between gene expression patterns of each prioritized gene. Blue squares indicate negative correlation. Red squares indicate positive correlation. Both are shaded on a gradient scale according to the Z score of the correlation. A dendrogram computed with Ward distances between the correlations is shown on top of the heatmap. Branches of the dendrogram are coloured differently to mark separate clusters. (**B**) Results of the REACTOME gene set enrichment analysis of the genes belonging to each of the clusters identified in (A). Colour key denotes the significance (−log 10 multiple testing adjusted *p* value) of each biological pathway. (**C**) Heatmaps depicting the scaled expression of prioritized genes belonging to the four clusters identified in (A) in three available RNA-seq datasets: intestinal biopsies from controls (CTR, n=5 samples) or CeD patients (CeD, n=6 samples); CD8^+^ TCRαβ intraepithelial cytotoxic lymphocytes (IE-CTLs) unstimulated or treated with IL-21, IL-15 or IFNβ for 3 hours (n=8 samples per condition) and gsCD4^+^ T cells unstimulated or treated with anti-CD3 and anti-CD28 (aCD3) for 3 hours (n=22 samples per condition). Clustering was performed using the “average” method in hclust().

We could not identify a specific biological process linked to our first co-regulation cluster. However, genes such as *ULK3* (relevant for autophagy^50^) and *CSK (*relevant to T cell receptor (TCR) signaling^51^) are included in this co-regulation cluster. Our second cluster encompasses genes (e.g. *STAT1*, *CD274* and *IL12A*) implicated in interferon gamma (IFNγ) signalling and interleukin (IL)-6 signalling. Co-regulation cluster 3 contains genes (e.g. *CD28*, *CTLA4* and *ICOS)* associated with co-stimulation by CD28, a process that is essential for modulating T cell–activation. Finally, co-regulation cluster 4 contains chemokine (e.g. *CCR1*, *CCR2* and *CCR3*) and cytokine signalling genes (e.g. *IL2RA*, *IL21* and *IL18R1*) (**Fig. 2B**). The biological processes overrepresented in these co-regulation clusters are essential for the activation and function of the adaptive and innate immune system, which confirms and extends previous findings that implicate both arms of the immune system in CeD disease pathogenesis. Approximately 10% of the prioritized genes are long non-coding RNAs (lncRNAs) rather than protein-coding genes (**Supplementary Table 1**). Although little is known about the function of lncRNAs, their co-regulation pattern with the genes in clusters 2 and 4 suggests that they may be associated with cytokine/chemokine signalling (**Fig. 2A, B**). Moreover, by using Genenetwork^34^, we found that the lncRNAs *RP3-395M20.9*, *AC007278.2* and *AC104820.2* may be involved in tumour necrosis factor (TNF) signalling, neutrophil degranulation and chemokine receptor signalling, respectively, implying a role for these uncharacterized lncRNAs in immune regulation in CeD.

### CeD candidate genes operate in immune and intestinal epithelial cells

To complement our REACTOME gene set enrichment analysis and dig deeper into the biological processes and cell types in which the prioritized genes may act, we analysed their expression profiles in available RNA-seq datasets from disease-relevant cell types including 1) small intestinal biopsies of active CeD patients and healthy controls, 2) intra-epithelial cytotoxic lymphocytes (IE-CTLs) stimulated with disease-relevant cytokines IL-21, IL-15 and IFNβ, and 3) gluten specific CD4^+^ T cells (gsCD4^+^ T cells) stimulated with antiCD3-antiCD28, which mimics the disease-specific response to gluten peptides (**Fig. 2C**) (differentially expressed genes for each dataset are available in **Supplementary Table 6**). We observed that the genes grouped in co-regulation clusters 1 and 2 are highly expressed in small intestinal biopsies and IE-CTLs, which is in line with the IFNγ pathway enrichment seen in co-regulation cluster 2 (**Fig. 2B**). IFNγ is mainly produced by gsCD4^+^ T cells and IE-CTLs and is known to disrupt the integrity of the intestinal epithelial cells in CeD-associated villous atrophy^52–54^. Within this cluster we also found genes specific to antigen-presenting cells (B cells, monocytes and dendritic cells) and epithelial cells such as *IL12A* and *COLCA1*, which are most expressed in small intestinal biopsies (**Fig. 2C**). The genes in co-regulation clusters 3 and 4 are highly expressed in gsCD4^+^T cells, especially after stimulation with antiCD3-antiCD28, indicating that these prioritized genes may be biologically relevant in the immediate T cell receptor response to gluten ingestion.

The gene expression pattern of the prioritized genes, when combined with information from our literature search, suggests that these genes may control general biological processes (e.g. apoptosis, gene regulation and cytoskeleton remodelling) as well as specific immune functions (e.g. cell adhesion, cell differentiation and TCR signalling) in diverse cell types (e.g. T cells, neutrophils, B cells, monocytes, epithelial cells) **(Fig. 3** and **Supplementary Table 7**). The non-HLA genetic loci associated to CeD thus seem to affect a complex network of cells and biological processes.

**Fig. 3.**
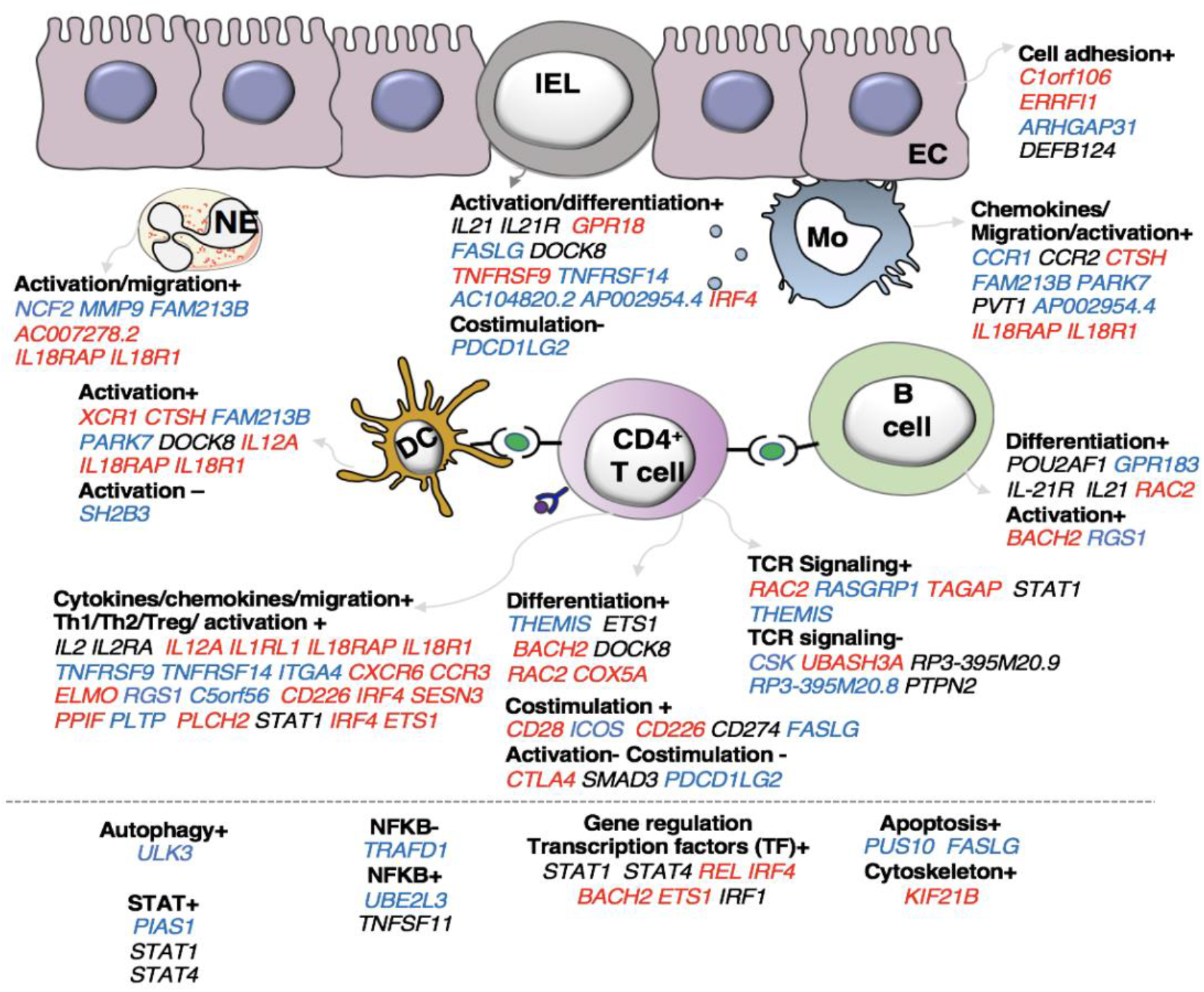
CeD candidate genes operate in immune cells and intestinal epithelial cells. Functions and cell types highlighted by the prioritized genes, according to our literature review (see **Methods**) (n=118 genes, for 37 genes neither a function nor a specific cell type on which the gene may operate could be specified). All genes contributing to a specific function are listed under the sub-heading and coloured according to the change that leads to increased CeD risk: increased expression (red), decreased expression (blue), or undefined (black). The symbols + or – denote if a biological process is thought to be induced or repressed by the gene, respectively, according to literature.

### Mediation analysis uncovers *TRAFD1* as a major *trans*-eQTL regulator

To further understand the potential regulatory function of the prioritized genes, we identified downstream regulatory effects by performing a *trans*-mediation analysis using a two-step approach (**Methods**) (**Supplementary Fig. 1A**). We first considered all genes with a *trans*-eQTL (p < 5×10^−8^) located in any of the 58 CeD-associated regions, then performed a mediation analysis by re-assessing the *trans*-eQTL effect after adjusting the expression levels for the expression of the prioritized gene(s) in the same locus (**Fig. 4A**).

**Fig. 4.**
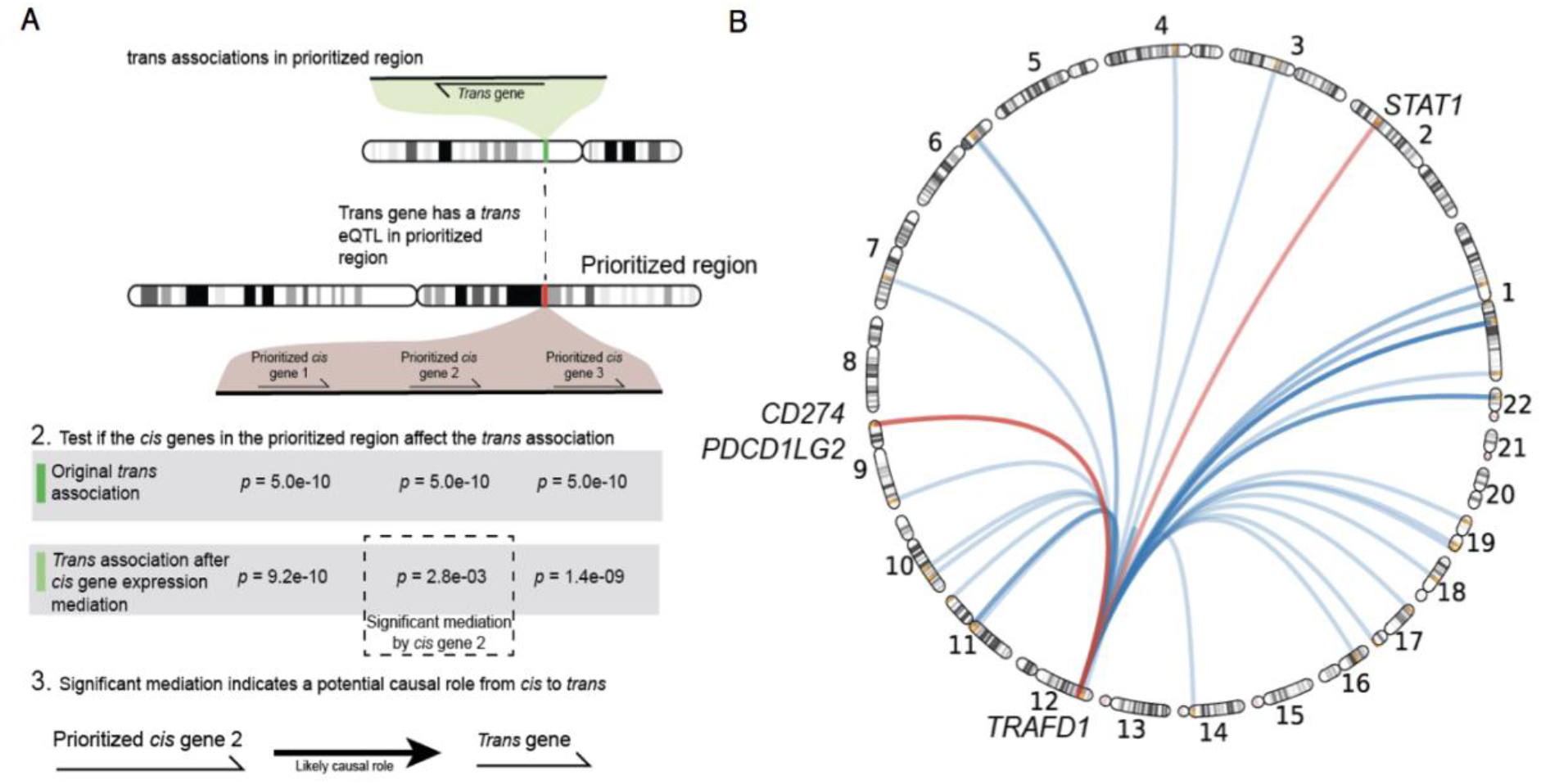
Mediation analysis uncovers *TRAFD1* as a major *trans*-eQTL regulator. (**A**) Workflow illustrating the main steps to identify *trans*-eQTL genes mediated by our *cis*-prioritized genes. First, we identified *trans-*eQTLs and *trans* genes that have a significant association (p < 5×10^−8^) in our prioritized regions. Then, for every *cis* prioritized gene in the CeD-associated region, a mediation analysis was performed to determine if the *cis* gene expression explains the *trans*-eQTL effect. (**B**) Circular ideogram depicting the mediating effect of *TRAFD1* on 41 trans genes. Three of the 41 *trans*-mediated genes were also prioritized by our *cis*-eQTL analysis (red).

Of the 497 possible prioritized gene–*trans*-eQTL gene combinations, we found 172 that exhibited significant mediation effects. These combinations map to 13 associated regions and represent 21 unique mediating *cis*-eQTL genes and 79 unique mediated *trans-eQTL* genes (**Supplementary Table 8**). Among all the associated regions, the CeD-associated region on chromosome 12 contained the largest number of both *cis*-mediating genes (N=5) and *trans*-mediated genes (N=60). In this region, *TRAFD1* mediated more *trans* genes than all of the other regional *cis*-regulators and also had the highest mediation impact (average Z-score difference in effect size between mediated and unmediated analysis = 2.79) (**Methods**) (**Supplementary Table 8**) (**Supplementary Fig. 1B**). Of note, the top eQTL variant of *TRAFD1* is a missense variant in the nearby gene *SH2B3.* This missense variant has been associated to a number of complex traits, including blood cell types and platelets, and autoimmune diseases^55,56^. However, we found that cell-type composition did not affect the eQTL-association of *TRAFD1* in our cohort (*p* > 0.044 for 24 different cell-type traits) (**Methods**) (**Supplementary Table 9**). To ensure that the mediated *trans* genes of *TRAFD1* were not mediated by *SH2B3*, we corrected *TRAFD1* expression levels for *SH2B3* and re-ran the mediation analysis. Here we found that the mediating effect of *TRAFD1* was still significant for all 41 genes found initially and that the median Z-score difference between mediated and unmediated was higher than that of *SH2B3*, although it was slightly attenuated compared to the original *TRAFD1* signal (**Supplementary Table 10**) (**Supplementary Figure 1B**). Based on these results, we conclude that *TRAFD1* is a master regulator of gene expression changes in the associated region (**Fig. 4B**) (**Supplementary Table 10**).

Strikingly, three of the *TRAFD1 trans*-mediated genes – *STAT1*, *CD274* and *PDCD1LG2* – are also prioritized *cis*-genes in their respective loci (**Fig. 4B**). These results suggest that the *trans*-mediated *TRAFD1*-effects may have an additional additive effect in these CeD-associated loci.

*TRAFD1* is a poorly characterized gene that has been suggested to act as a negative regulator of the NFκB pathway^57^. To further elucidate the biological processes in which the 41 *TRAFD1 trans*-mediated genes could be involved, we performed a REACTOME 2016 gene set enrichment analysis (**Supplementary Table 11**). Here we found that IFNγ signalling, cytokine signalling and major histocompatibility complex class I (MHCI) antigen processing / presentation are strongly enriched pathways, which points to a role for *TRAFD1* and *TRAFD1 trans*-mediated genes in antigen presentation and immune response (**Fig. 5A**).

**Fig. 5.**
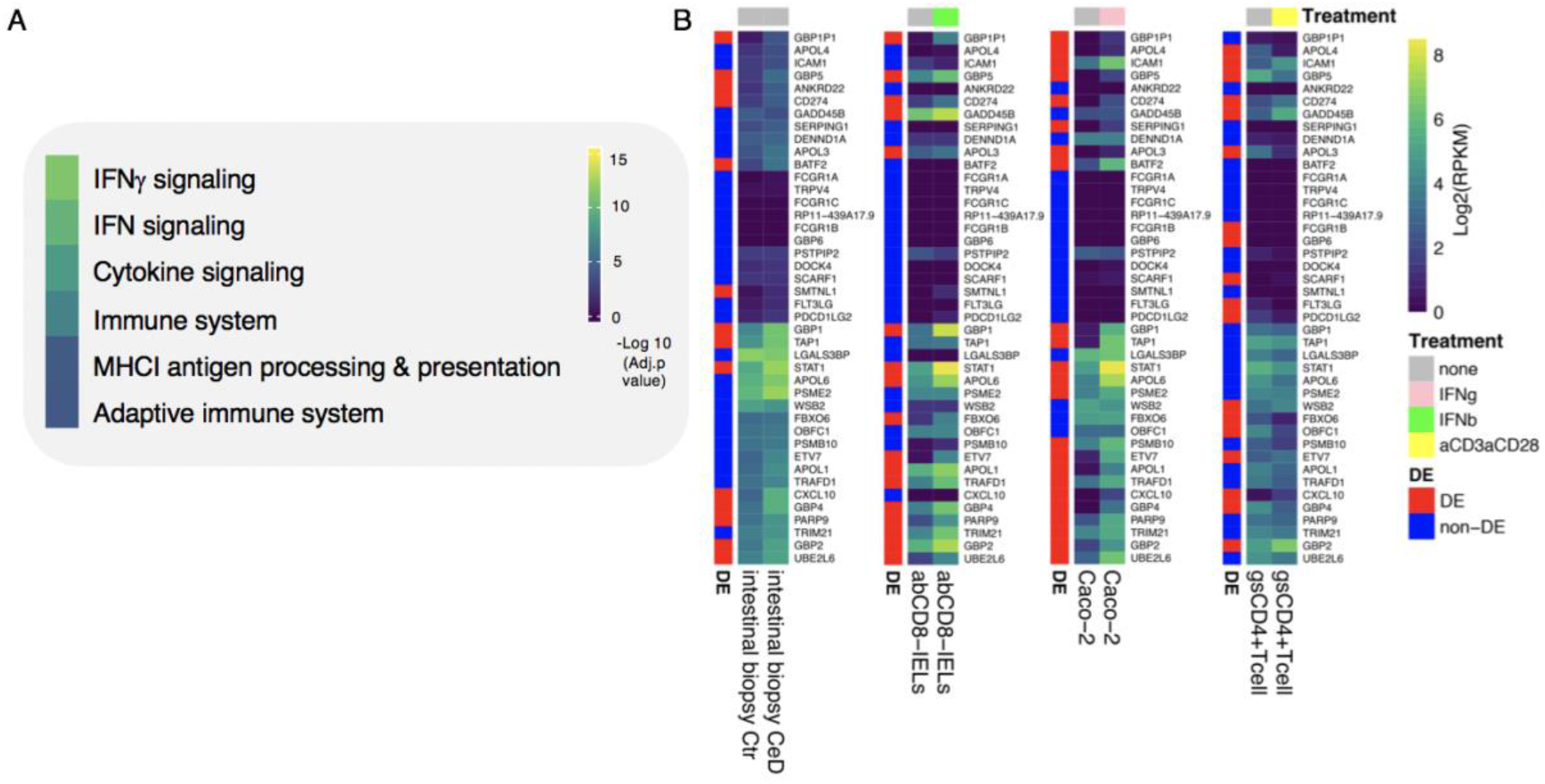
*TRAFD1* is a regulator of IFNγ signalling genes. (**A**) Results of the REACTOME gene set enrichment analysis of *TRAFD1*-mediated genes (n=41 genes). Colour code denotes the significance (−log 10 adjusted *p* value) of each biological pathway. (**B**) Unscaled heatmaps depicting the expression of these genes in RNA-seq datasets from different cell types: whole biopsies from controls (Ctr, n=5 samples) of CeD patients (CeD, n=6 samples); intraepithelial cytotoxic lymphocytes (IE-CTLs) unstimulated or treated with IFNβ for 3 hours (n=8 samples per condition); and Caco-2 cells untreated or stimulated with IFNγ for 3 hours (n=8 samples per condition). Red indicates that a gene is differentially expressed (DE), blue indicates that a gene is not differentially expressed (non-DE) (FDR<0.01 and |log2(RPKM)>1|). Grey (none or unstimulated), pink (IFNγ), green (IFNβ) and yellow (antiCD3/antiCD28) colours indicate the type of stimulation (treatment).

By looking into RNA-seq datasets from disease-relevant cell types, we noted that most *TRAFD1 trans*-mediated genes are upregulated in biopsies from patients with active CeD, and these genes include *STAT1*, *CXCL10* and *TAP1*, which are essential for IFN response^58^, chemotaxis^59^ and antigen processing^60^, respectively (**Fig. 5B**). Moreover, most *TRAFD1 trans-*mediated genes exhibit an increase in expression in response to IFNγ in intestinal epithelial cells (Caco-2) or IFNβ in IE-CTLs (**Fig. 5B**). In contrast, antiCD3-antiCD28 stimulation in gsCD4^+^T cells resulted in both up- and downregulation of the *TRAFD1 trans*-mediated genes, implying that *TRAFD1 trans*-mediated genes respond more strongly to IFN signalling (IFNγ or IFNβ) than to TCR activation by anti-CD3/anti-CD28. Indeed, the enrichment of the 41 *TRAFD1 trans*-mediated genes in significantly differentially expressed genes in biopsies, IE-CTLs, epithelial cells and gsCD4^+^Tcells was strongest in IE-CTLs and epithelial cells upon IFN signalling (**Supplementary Table 6**). Overall, our results suggest that *TRAFD1* and *TRAFD1 trans*-mediated genes modulate IFN signalling upon antigen presentation, possibly via regulation of NFκB, in CeD pathology.

### *TRAFD1* KD affects immune-activation genes

We performed a siRNA KD experiment on *TRAFD1* to gain more insights into the biological function of this gene and to independently validate the *TRAFD1 trans*-mediated genes. We also evaluated the transcriptional changes of knocking down *TRAFD1* in the monocyte-like cell line THP-1 under resting conditions (unstimulated) or in the presence of LPS, a known inducer of the NFκB pathway^61^.

After siRNA treatment, we observed no significant differences in cell viability or proliferation among the controls (WT and SCR) and the KD treatment (**Supplementary Fig. 2A, B**). However, as expected for the KD cell line, we noted a significant reduction in the expression of *TRAFD1* compared to the controls in WB and qPCR analyses (**Supplementary Fig. 2C-E**). KD of *TRAFD1* was also confirmed in the RNAseq data, with *TRAFD1* expression levels reduced by 41% in unstimulated KD cells compared to unstimulated SCR cells (adjusted *p* = 0.004) and by 34% in LPS-stimulated KD cells compared to LPS-stimulated SCR cells (not significant) (**Supplementary Table 12**). The reduced KD effect upon LPS stimulation is consistent with our expectation that *TRAFD1* acts as negative regulator of the NFκB pathway, which is activated by several stimuli, including LPS^61^. Thus, the KD was successful and neither the transfection method nor a reduced expression of *TRAFD1* had a toxic effect (**Supplementary Fig. 2A-E**).

Next, we tested if the 41 *TRAFD1 trans*-mediated genes were more differentially expressed than expected after LPS stimulation (**Supplementary Fig. 3**). To disentangle differential expression from the co-expression inherently present in a gene expression dataset, we devised a permutation scheme that compared the control (WT vs. SCR) observations with the KD (SCR vs. KD) observations (see **Methods**). This scheme takes into account the co-expression of a gene set, as this co-expression is present in both the control and the experimental observation. After performing 1,000,000 permutations of 42 genes (41 *trans* mediated genes and *TRAFD1*) in the LPS-stimulated comparison, the median test statistic in the control observations was observed 54 times more often than in the KD observations (0.270% for WT-SCR vs. 0.005% for SCR-KD, **Supplementary Fig. 4**). This indicates that the 41 *trans*-mediated genes and *TRAFD1* are 54 times more differentially expressed than expected. We did not find increased differential expression of the same gene set in the unstimulated condition (1.120% for WT-SCR vs. 0.307% for SCR-KD, **Supplementary Fig. 4**), indicating that *TRAFD1* mainly regulates genes in an LPS-stimulated state.

To identify the role of *TRAFD1* in immune cells and processes, we compared gene expression changes in the unstimulated condition versus the LPS stimulated condition for each treatment (WT, SCR or KD) separately (**Supplementary Fig. 2F**). Differential expression analysis showed that 353 genes were uniquely upregulated and 330 genes uniquely downregulated after *TRAFD1* KD treatment (**Supplementary Fig. 2G, H**). We found no REACTOME gene set enrichment for these unique KD genes. We found 500 upregulated and 433 downregulated genes that were differentially expressed in all three treatments upon LPS stimulation (**Fig. 6A**, **Supplementary Fig. 2G, H**). Upregulation (or downregulation) after LPS stimulation was treatment-dependent, i.e. the differential expression identified was increased (or decreased) from WT to SCR to KD (**Fig. 6A**). We performed hierarchical clustering, this separated the two gene sets into two clusters: cluster 1 shows a decreased response of genes in the *TRAFD1* KD group (LPS cluster 1, **Fig. 6B**, **Supplementary Table 13**) and cluster 2 displays an increased expression in the *TRAFD1* KD cells under unstimulated conditions that persists after LPS stimulation (LPS cluster 2, **Fig. 6C**, **Supplementary Table 13**). REACTOME gene set enrichment analysis indicated that the genes in LPS cluster 1 are involved in immune-related processes (e.g. cytokine signalling, RIG/IMDA5 induction of IFN signalling and IFN signalling, **Fig. 6D**), whereas the genes in LPS cluster 2 are associated with the heat shock response, which has been shown to be activated as a consequence of immune activation or immune response to stress^62^ (**Fig. 6E**, **Supplementary Table 13**). Together, these results suggest that *TRAFD1* is a regulator of immune activation and inflammation.

**Fig 6.**
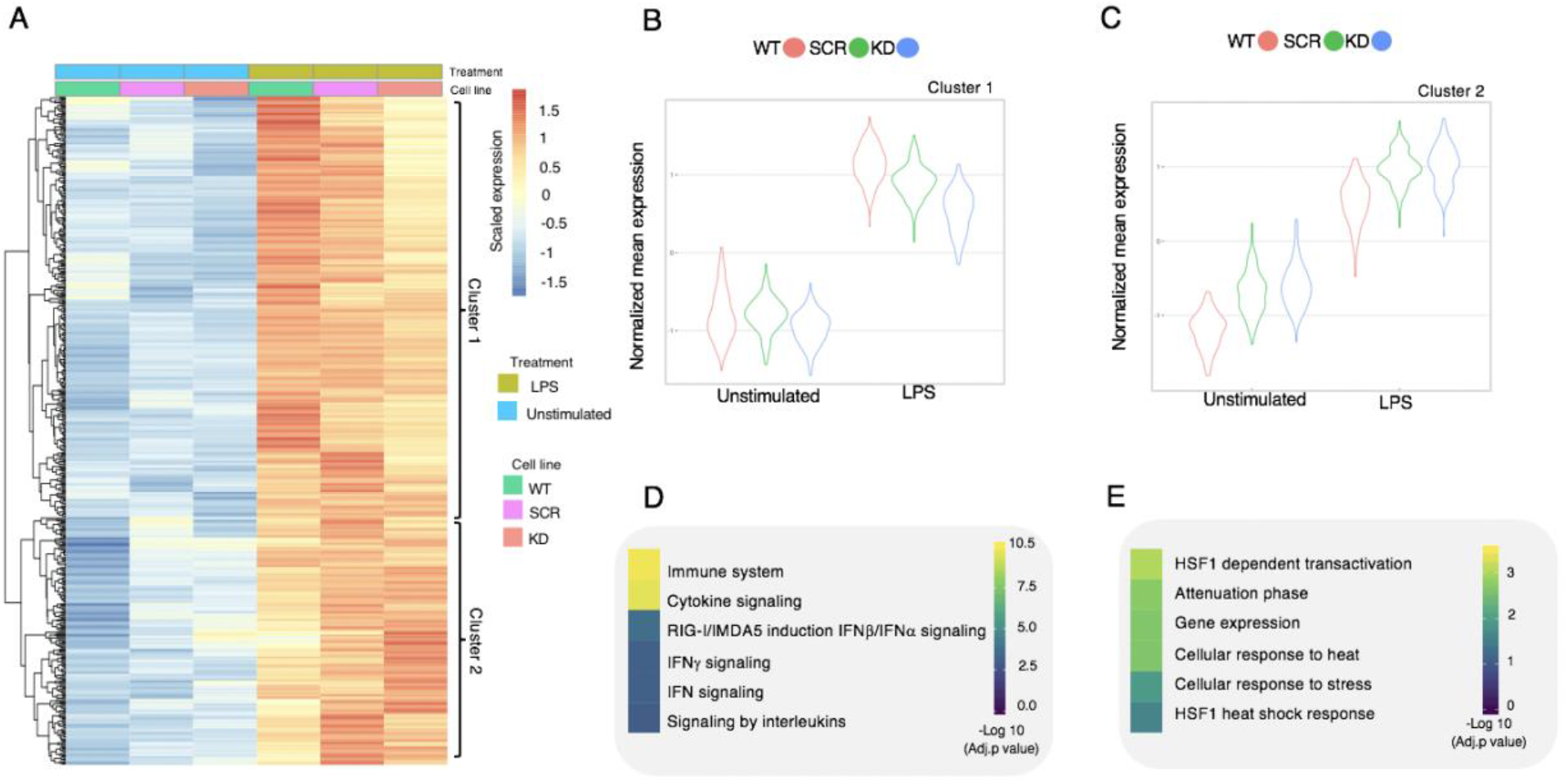
*TRAFD1* knockdown affects immune activation and stress-related genes. (**A**) Heatmap showing the expression profile of the 500 shared DEGs identified in the knockdown experiments (see **Methods** and **Supplementary Fig. 2 F**, **G**). A dendrogram on the left of the heatmap depicts the strength of similarities based on Ward distance. (**B, C**) Violin plots showing the normalized gene expression of the genes belonging to the first and second cluster of DEGs identified in (A) in THP-1 cells under different experimental conditions (WT=untransfected, SCR=non-targeting siRNA, KD=siRNA targeting *TRAFD1*) and stimulations (LPS=lipopolysaccharide). (**D, E**) Results of REACTOME gene set enrichment analysis of the genes within the first (D) and second cluster (E). Significance (−log 10 adjusted *p* value) of each biological pathway is indicated by the colour key.

## Discussion

In the present study we aimed to identify CeD candidate genes using four *in silico* methods (MR-IVW, COLOC, LD overlap and DEPICT) and whole blood transcriptomics data from a population-based cohort. While previous studies have used at least one of these methods^3,5,6,11^, to our knowledge this is the first effort that integrates the four different statistical approaches. This systematic prioritization approach resulted in 118 prioritized causal genes, including 26 that are direct targets of an approved drug or of drugs under development for other complex diseases, including autoimmune diseases. The co-expression pattern within a large RNA-seq dataset from blood^34^ suggests these genes are involved in cytokine signalling in innate and adaptive cells as well as in T cell activation pathways. We also identified *TRAFD1* to be *trans*-regulator of 41 genes, with a strong enrichment in IFNγ signalling and MHC I antigen processing/presentation pathways, which are pivotal for the disease pathogenesis.

After clustering our *cis*-eQTL prioritized genes on shared co-regulation, we identified a cluster of genes involved in T cell activation and co-stimulation (cluster 3), highlighting a key role for T cell activation in the pathogenesis of CeD^63^. Within this co-regulation cluster we found the *THEMIS*, *IL2, CD28*, *CTLA4* and *UBASH3A* genes (**Fig. 2**), whose functions include T cell differentiation and activation and the TCR macromolecular complex.

Another co-regulation cluster (cluster 4, **Fig.2**) grouped prioritized genes involved in cytokine and chemokine signalling events that affect the microenvironment during inflammation in the intestinal mucosa. For example, this group included *CCR1* and *CCR2*, which control the activation and recruitment of inflammatory cells such as monocytes, dendritic cells and neutrophils^64^. *IL21* was also included in this co-regulation cluster. This gene encodes IL-21, providing proliferation and survival signals to B cells^65^, which in turn produce the autoantibodies detected in CeD and could act as antigen-presenting cells for gsT cells, thus enhancing the inflammatory response^66^. Next to chemokine receptors, this co-regulation cluster contains cytokine receptor genes, e.g. *IL18RAP* and *IL18R1*. Which encode the IL-18 receptor, that is expressed in intestinal epithelial cells and mediates IL-18 controlled intestinal barrier integrity and immunity^67,68^. This cluster also contains transcription factors genes, e.g. *IRF4*, *ETS1* and *REL*. *IRF4* and *ETS1* are essential for T helper 1 (Th1) differentiation^69,70^. Interestingly, gsCD4^+^ T cells exhibit a Th1 profile, that predominantly produce IFNγ, a cytokine that affects the integrity of the intestinal epithelial cells contributing to villous atrophy^52–54^. *REL*, that is also contained in cluster 4, is a key regulator of NFκB signalling pathway, a major mediator of inflammation^71^, which is in line with the novel genetic association reported between NFκB and CeD by Ricano-Ponce et al.^5^ Moreover, CeD patients show a persistent activation of the NFκB pathway in the intestinal mucosa^72^ as well as a significant increase in the methylation level of 8 genes that belong to this pathway^73^. Thus, these results indicate that CeD patients present with a major defect in the NFκB signalling complex.

For practical reasons, most prioritization studies have been focused on incorporating *cis*-eQTLs^74^ and have mostly ignored trans-eQTLs, thus potentially missing long-distance co-regulated interactions^74^. In our study, we took advantage of a large transcriptomics cohort to run a *trans* mediation analysis for CeD loci. One of the most remarkable findings of this approach was that 41 *trans*-mediated genes were found to be controlled by a single gene: *TRAFD1*. These 41 genes are enriched for IFNγ and MHC I antigen processing/presentation signalling pathways. Interestingly, gsCD4^+^ T cells exhibit a Th1 profile and produce a large amount of IFNγ, one of the most predominant cytokines in CeD^54^. Some of the most striking effects of IFNγ include induction of apoptosis in intestinal epithelial cells, alteration of intestinal permeability and activation of monocytes and dendritic cells, which may act as antigen-presenting cells for gsCD4^+^ T cells^75^.

*TRAFD1* is thought to be a regulator of the NFκB signalling pathway^57^, suggesting that CeD-risk SNPs may modulate the NFκB complex via both *cis* and *trans* regulatory mechanisms. Our results also point to a role for *TRAFD1* in response to IFNγ; however, IFNγ does not typically activate NFκB signaling^76^ and the *IFNG* locus is not associated with CeD^7^. Thus, *TRAFD1* may activate the production of other cytokines, which in turn activate the NFκB complex.

IE-CTLs, which are the effector cells in CeD, have not thus far been genetically associated with the disease. However, given that MHC-I antigen presentation presentation/processing are essential for IEL activation and the striking activation of the 41 *trans* mediated genes in IE-CTLs upon IFN stimulation, we propose that the IE-CTLs are also genetically linked to the disease through the action of *TRAFD1*.

Despite the approaches implemented in our study to uncover the novel gene interactions and biological pathways that may underlie the disease, a major drawback is the limited genome coverage of the CeD summary statistics used in this study. These were derived from a GWAS that used the Immunochip platform, a genotyping platform that only measures genotypes in regions known to be associated with immune function. We thus acknowledge that our current interpretation of CeD loci is biased toward immune-related mechanisms. Only when comprehensive whole-genome CeD association analyses have been performed will we have an unbiased understanding of the disease pathophysiology.

In our gene prioritization we observed that the different statistical gene prioritization methods applied to our data prioritized unique and jointly prioritized genes. Therefore, we recommend that investigators incorporate multiple methodologically orthogonal gene prioritization methods to identify a more comprehensive set of causal genes for a given disease. Here, we use two different (orthogonal) expression datasets (BIOS and DEPICT) and three prioritization methods using the same underlying data: MR-IVW, LD-overlap and COLOC. While we believe that the genes prioritized in this study represent robustly prioritized genes for CeD, it is difficult to validate if all the prioritized genes are truly causal based on statistical methodology alone. Functional validation of these genes in disease context is needed to rule out false positives.

The functional validation of *TRAFD1* in the siRNA KD experiment in THP-1 cells does establish that this gene regulates the *trans*-mediated network identified by our eQTL and statistical analysis. Still, the effects of the SCR control and the transfection itself may have obscured some specific *TRAFD1*-mediated effects. Moreover, the CeD-associated effects of *TRAFD1* may not be most pronounced in monocytes or upon LPS-stimulation. Indeed, context- and cell-type-specific effects of CeD-associated genetic variation may hamper the identification of the downstream effects of the prioritized *cis-* and *trans*-genes.

In conclusion, this study provides a framework for predicting candidate genes and their function using a systematic *in silico* approach that could be extended to other complex diseases. Using this approach, we not only confirmed previous association between adaptive cells (gsCD4^+^ T cells and B cells) and CeD, we also unveiled a link between specific genes that may contribute to the disease via innate immune cells, epithelial cells and IE-CTLs. Finally, we found a gene network controlled by *TRAFD1* that is part of two major pathways of immune activation, IFNγ signalling and MHC I antigen processing.

## Glossary

Underlined words are definitions that have been explained in the preceding lines

eQTL: expression quantitative trait locus, a location on the genome that is statistically associated to changes in gene expression.
*cis*-eQTL: an eQTL located in the same locus of the gene that is being interrogated (within 1.5Mb from gene transcript start or end).
*trans*-eQTL: an eQTL that is not physically close to the gene that is being interrogated (>1.5Mb from transcript start/end or on a different chromosome).
*cis*-eQTL gene: a gene that is associated with a change in expression as a consequence of a *cis*-eQTL.
*trans*-eQTL gene: a gene that is associated with a change in expression as a consequence of a *trans*-eQTL.
CeD: celiac disease
CeD-associated region: a genomic region that is associated to CeD based on results from genome-wide association studies on CeD.
Prioritized gene: a gene prioritized as being potentially causal for CeD according to the four statistical methods depicted in Figure 1A-B. In this study, prioritized genes are always within the CeD-associated regions.
Mediating *cis* gene: a prioritized gene that is statistically responsible for the change in expression of a *trans*-eQTL gene. Of note, while the *trans*-eQTL is located in the same CeD-associated region of the mediating *cis*-gene, the mediated *trans-*gene is not.
Mediated *trans* gene: a gene located outside CeD-associated regions that is statistically mediated by a mediating *cis* gene located in the same region of the corresponding *trans*-eQTL.

## Acknowledgements

We are very grateful for the altruistic donation of biological materials by the study participants, without them this study would not be possible. In addition, we thank the UMCG Genomics Coordination center, and the UG Center for Information Technology, and their sponsors BBMRI-NL & TarGet, for storage and computing infrastructure. We thank BBMRI-NL for providing the transcriptome and genotyped data for the BIOS cohort. We would like to thank Prof. Bana Jabri for providing IE-CTL cell lines, Prof. Morris Swertz for data storage and cluster facilities and Kate McIntyre for editing the manuscript. This work was supported by an ERC Advanced grant [FP/2007-2013/ERC grant 2012-322698] and an NWO Spinoza prize grant [NWO SPI 92-266] to C.W. I.J is supported by a Rosalind Franklin Fellowship from the University of Groningen and an NWO VIDI grant [016.171.047].

## Data Availability

Summary statistics of the CeD GWAS are available from the European Genome-Phenome Archive (https://www.ebi.ac.uk/ega/studies/EGAS00001003805) under accession number EGAS00001003805. The individual-level data of the BIOS cohort is available upon request from https://www.bbmri.nl/acquisition-use-analyze/bios.

## Supplementary figure legends

**Supplementary Fig. 1 Mediation effect on *trans* genes for all prioritized genes in the *TRAFD1* region on chromosome 12**. (**A**) Three boxes with the eQTL association curves of *TRAFD1*, *SERPING1* and *SERPING1* after mediation with *TRAFD1*. (**B**) Scatter plot indicating the absolute Z difference between unmediated and mediated *trans* associations upon mediation (y axis) by all mediating *cis* genes in the *TRAFD1* region shown on the x axis as well as when correcting *TRAFD1* expression for the expression of *SH2B3* (‘*TRAFD1 – SH2B3’*).

**Supplementary Fig. 2 *TRAFD1* knockdown validation.** Cell viability (**A**) and proliferation (**B**) of THP-1 cells that were left untransfected (WT) or transfected with non-targeting siRNA (SCR) or siRNA targeting *TRAFD1* (KD) for 72 hours. Protein and mRNA levels of *TRAFD1* were determined by WB (**C**, **D**) and qPCR (**E**). Bars indicate mean ± SEM. Data are representative of three different experiments. Statistical differences were calculated with a one-sided t-test by using the SCR as 100% reference. p-value ≤ 0.0001 (****). (**F**) The differential expression analysis approach. Here we compared the gene expression between unstimulated samples and their respective LPS-stimulated samples to identify DEGs that respond to stimulation ((|log2 FC|) >1 and FDR ≤ 0.01). We then identified unique or shared DEGs responding to the stimulation between treatments (WT, SCR or KD), which are shown in two separate Venn diagrams: one for upregulated genes (**G**) and one for downregulated genes (**H**).

**Supplementary Fig. 3 Expression pattern of *TRAFD1*-mediated genes upon *TRAFD1* knockdown.** Heatmap showing the pattern of gene expression of *TRAFD1* and of the 41 genes it mediates, scaled by row (see details in **Methods** and **Fig. 5**). Expression is shown in different treatments and stimulations as indicated by coloured bars on top of the heatmap.

**Supplementary Fig. 4 DEGs upon *TRAFD1* knockdown are enriched in *TRAFD1*-mediated genes.** Here we compare the differential expression of 42 genes found in the *trans* mediation analysis of *TRAFD1* (41 *trans*-mediated genes and *TRAFD1*) with the differential expression of 42 other randomly chosen genes. The histograms (blue) show the distribution of the median absolute T statistic of DEseq of 42 randomly chosen genes, when 1,000,000 sets of genes are randomly chosen, compared to the observed value for the 42 genes that are from the *trans*-mediation analysis (red horizontal line). We compare the results of the control experiment (WT-SCR) in panels **A** and **C** with the results of the knockdown experiment (SCR-KD) in panels **B** and **D.** The fold differences between the control experiments and the knockdown experiments show how much more than expected the 42 genes are differentially expressed in the knockdown compared to the control.

## Supplementary table Legends

**Supplementary Table 1. Prioritization of genes likely causal for celiac disease (CeD).** This table contains all the genes in the prioritized CeD regions and their evidence for being causal to CeD. One gene per row is shown. Columns (in order): the human build 37 coordinates of the CeD region in which the gene is located (**region**); the gene name according to the ENSEMBL GENES 96 database (human build 37) (**gene_name**); the ENSEMBL gene identifier (**ensembl_id**); the most likely effect direction (determined as described in **Methods**) (**most_likely_direction**); number of independent eQTL variants found for the gene (**n_eqtl_effects**); the effect size (**MR_ivw_effect**) and *p* value (**MR_ivw_p_value**) of the MR-IVW test; the summary of LD overlap (**ld_overlap_summary**), with either the top eQTL variant (‘top_snp’) or an independent eQTL variant (‘cojo_snp’) with the r^2^ linkage disequilibrium between the eQTL SNP and the CeD top variant; the coloc posterior probability of causal variants being shared (**coloc_h4**); if the gene passes DEPICTs own false discovery thresholds (**depict_fdr_pass**); and the lines of evidence that are significant compared to the lines of evidence that are available for a gene (**lines_of_evidence**). Bold fields in any of the columns indicate that the prioritization method is significant according to our thresholds.

**Supplementary Table 2. Sensitivity analyses for genes selected by the IVW-MR method.** In this table, genes with a significant MR-IVW effect are tested for heterogeneity using the Q test statistic and the MR-weighted median results as sensitivity analysis of all significant MR results. Each row contains the following information: the human build 37 coordinates of the CeD region in which the gene is located (**region**); the gene name according to the ENSEMBL GENES 96 database (human build 37) (**gene_name**); the ENSEMBL gene identifier (**ensembl_id**); the most likely direction of the effect (determined as described in **Methods**) (**most_likely_direction**); the number of independent eQTL variants found (**n_eqtl_effects**); the effect size (**MR_ivw_effect**) and *p* value (**MR_ivw_p_value**) of the MR-IVW test; the heterogeneity *p* value of the MR-IVW test using Cochran’s Q statistic (**MR_heterogeneity_p_value**); the weighted median effect estimate (**MR_WM_beta**) and its associated *p* value (**MR_WM_p**); the MR effect estimate after removal of potential outliers (**MR_Q_beta**); its associated *p* value (**MR_Q_p**); the remaining variants after outlier removal (**MR_Q_ivs**) and the heterogeneity estimate (**MR_Q_heterogeneity**).

**Supplementary Table 3. Druggability information for prioritized genes.** This table contains all the prioritized *cis* genes in the CeD regions that are existing drug targets according to two different databases (DrugBank v5.1.4, and Finan et al.^39^). One gene per row is shown. Columns indicate (in order): the human build 37 coordinates of the CeD region in which the gene is located (**region**); the gene name according to the ENSEMBL GENES 96 database (human build 37) (**gene_name**); the ENSEMBL gene identifier (**ensembl_id**); the most likely effect direction (determined as described in **Methods**) (**most_likely_direction**); the number of independent eQTL variants found for the gene (**n_eqtl_effects**); the effect size (**MR_ivw_effect**) and *p* value (**MR_ivw_p_value**) of the MR-IVW test; the summary of LD overlap (**ld_overlap_summary**) with either the top eQTL variant (‘top_snp’) or an independent eQTL variant (‘cojo_snp’) with the r^2^ linkage disequilibrium between the eQTL SNP and the CeD top variant; the coloc posterior probability of causal variants being shared (**coloc_h4**); if the gene passes DEPICT’s own false discovery thresholds (**depict_fdr_pass**); the lines of evidence that are significant compared to the lines of evidence that are available for a gene (**lines_of_evidence**); the druggability tier based on Finan et al.^39^, with lower tiers making it more likely that the gene is druggable^39^, (**druggable_tier**), and if the gene is a drug bank drug target (**drug_bank_drug_target**). Bold fields in any of the columns indicate that the prioritization method is significant according to our thresholds.

**Supplementary Table 4. Cluster assignments for the prioritized genes.** The 118 prioritized genes were assigned to a cluster based on a guilt-by-association co-regulation approach to find shared biological mechanisms. For each gene that was prioritized (**ensembl_id** and **gene_name**), a cluster membership is given (**cluster_membership**).

**Supplementary Table 5. Significant REACTOME 2016 enrichment of *cis* prioritized genes in each co-regulation cluster**. Results from the enrichr API using the gene clusters of **Supplementary Table 4** as query. Columns indicate: the enrichment background (**background**); the enrichment term in the background (**term_name**); the non-corrected *p* value of enrichment for this term (**p_value)** and *Z* score (**Z_score**); enrichr combined score (**combined_score**); *cis*-prioritized genes found in each the term (**overlapping_genes**) and the multiple testing corrected *p* value (**adjusted_p_value**). Each tab of the excel file contain the gene set enrichment for each cluster as defined in **Supplementary Table 4**.

**Supplementary Table 6. Results of DE analyses from all cell-type- and context-specific data available for this study (datasets).** This table lists all results for the DE analyses (Significant DE genes are defined as padj < 0.05 and log2 fold change > |1|) and a summary report of the overlap with *TRAFD1 trans*-mediated genes (**overlap with trans genes+** *TRAFD1*) and relative enrichment. The DE gene lists (padj < 0.05 and log2 fold change > |1|) for each dataset are given in individual sheets. In the sheet “enrichment”, columns upregulated and downregulated indicate if the *trans*-mediated genes are up- or downregulated under stimulated conditions compared to control conditions in each dataset. Enrichment of all the trans mediated genes in the DE genes was determined using a Fisher’s exact test and the enrichment *p* value is shown in the column enrichment p-val.

**Supplementary Table 7. Functions attributable to the prioritized genes, according to our literature review (see Methods and Fig. 3).** Columns describe (in order): gene name (**gene_name**); ensemble ID (**ensembl_id**); the change that leads to increased CeD risk, i.e. increased expression (+), decreased expression (−), or undefined (?) (**direction**); attributable function based in literature (**potential_function**); and literature or web-based sources (**source_1** and **source_2**). Web-based sources include Gene cards (https://www.genecards.org/) and Genenetwork (https://www.genenetwork.nl/).

**Supplementary Table 8. All the significant *trans*-mediated genes from our *cis* prioritization.** Each row contains a *cis–trans* gene pair described with both the ensembl id and hgnc gene name (**cis_ensembl_id**), (**cis_gene_name**), (**trans_ensembl_id**) and (**trans_gene_name**). Mediation effect and significance are shown using the Z score of the unmediated versus the mediated estimate (using the original unmediated standard error) (**z_score_difference**) and the mediation *p* value of the test defined by Friedman and Schatzkin (**mediation_p**).

**Supplementary Table 9. Cell type mediation analysis.** We calculated to what extent cell types counted in the BIOS cohort affect the most highly associated *TRAFD1* eQTL variant. Columns show (in order): the specific cell type measurements or mediator (**mediator**); the effect size after mediation by the cell type (**mediated_beta**); the original effect size (**unmediated_beta**); difference in effect sizes between mediated and unmediated (**beta_difference**); the standard error mediation effect size (**se**); the t-statistic of the beta differences (**t_statistic**); a *p* value of the Friedman and Schatzkin test statistic (**p_value**); the Pearson correlation coefficient between *TRAFD1* and the cell type proportion (**correlation**); and the number of observations in the BIOS cohort (**n_observations**). If a mediator has a “_Perc” suffix, the cell type counts were converted into ratios. Cell type abbreviations: Baso: Basophil count, EOS: eosinophil count, HCT: haematocrit, HGB: haemoglobin, LUC: large unstained cell count, Lymph: lymphocyte count, MCH: mean corpuscular haemoglobin, MCHC: mean corpuscular haemoglobin concentration, MCV: mean corpuscular volume, Mono: Monocyte count, MPV: mean platelet volume, Neut: Neutrophil count, PLT: platelets count, RBC: red blood cell count, RDW: red blood cell distribution width, WBC: white blood cell count.

**Supplementary Table 10. Mediation results when correcting *TRAFD1* expression for the nearby *SH2B3* expression**. Columns are: the ENSEMBL id (**ensemble_id**); the hgnc gene name (**gene_name**); and the mediation Z score difference (**Z_score_difference**), *p* value (**p_value**) and Pearson correlation (**correlation**) between the *trans*-eQTL top variant and the residual of *TRAFD1* expression, after correction for *SH2B3* expression.

**Supplementary Table 11. Significant REACTOME 2016 enrichment of *TRAFD1-* mediated genes.** Results from the enrichr API using the mediated *TRAFD1 trans* genes as query. Columns indicate: the enrichment background (**background**); the enrichment term in the background (**term_name**); the non-corrected *p* value of enrichment for this term (**p_value)** and *Z* score (**Z_score**); enrichr combined score (**combined_score**); *cis*-prioritized genes found in each the term (**overlapping_genes**) and the multiple-testing-corrected *p* value (**adjusted_p_value**).

**Supplementary Table 12. Differential expression results of the THP-1 experiments.** This table shows differential expression analysis of the THP-1 cells with 3 hr LPS treatment (LPS) or without LPS (Unstim) in wild type (WT), scrambled control siRNA (SCR) or TRAFD1 knock down conditions (KD). All conditions and treatments were performed in triplicate. Complete DESEQ2 results are shown for each possible comparison in each tab. For each gene the columns show: the ensembl id per gene (**ensembl_id**); the mean corrected expression of the gene (**baseMean**); the log2 fold change of the comparison (**log2FoldChange**); the standard error of this log2 fold change (**lfcSE**); a t-statistic of the log2foldchange (**stat**); the *p* value (**pvalue**); and the multiple testing adjusted *p* value (**padj**). The direction of the effect is always towards the second term in the tab name: if a log2 fold change is positive and the tab name is, for example, ‘WT_LPS_vs_SCR_LPS’, then the expression of the gene is increased in the SCR_LPS samples compared to the WT_LPS samples.

**Supplementary Table 13. Significant REACTOME 2016 enrichments in genes cells that are significantly upregulated by LPS in all treatments (WT, SCR and KD).** Reactome enrichment is shown for genes according in two groups: genes relatively downregulated in the *TRAFD1* knockdown experiment (REACTOME_enrichment_cluster1) and genes relatively upregulated in the TRAFD1 knockdown experiment (REACTOME_enrichment_cluster2). Results are shown from the enrichr API analysis using the genes in a cluster as query. Columns indicate: the enrichment background (**background**); the enrichment term in the background (**term_name**); the non-corrected *p* value of enrichment for this term (**p_value)** and *Z* score (**Z-score**); enrichr combined score (**combined_score**); *cis*-prioritized genes found in each the term (**overlapping_genes**) and the multiple testing corrected *p* value (**adjusted_p_value**).

