## Supplementary material for "Systematic prioritization of candidate genes in disease loci identifies *TRAFD1* as a master regulator of IFNγ signalling in celiac disease": Sup_Figures

### Slide 1
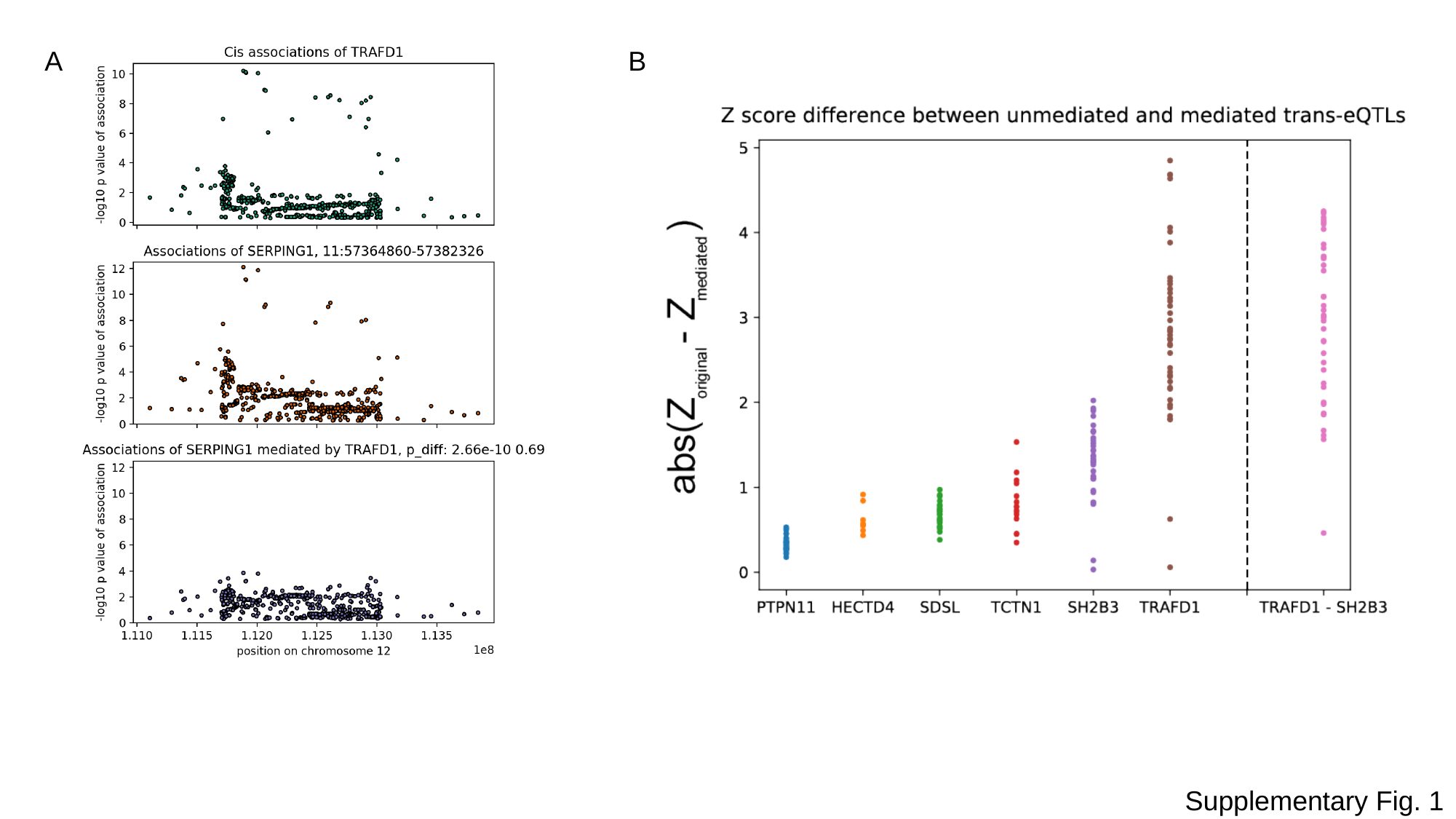

A
B
Supplementary Fig. 1

### Slide 2
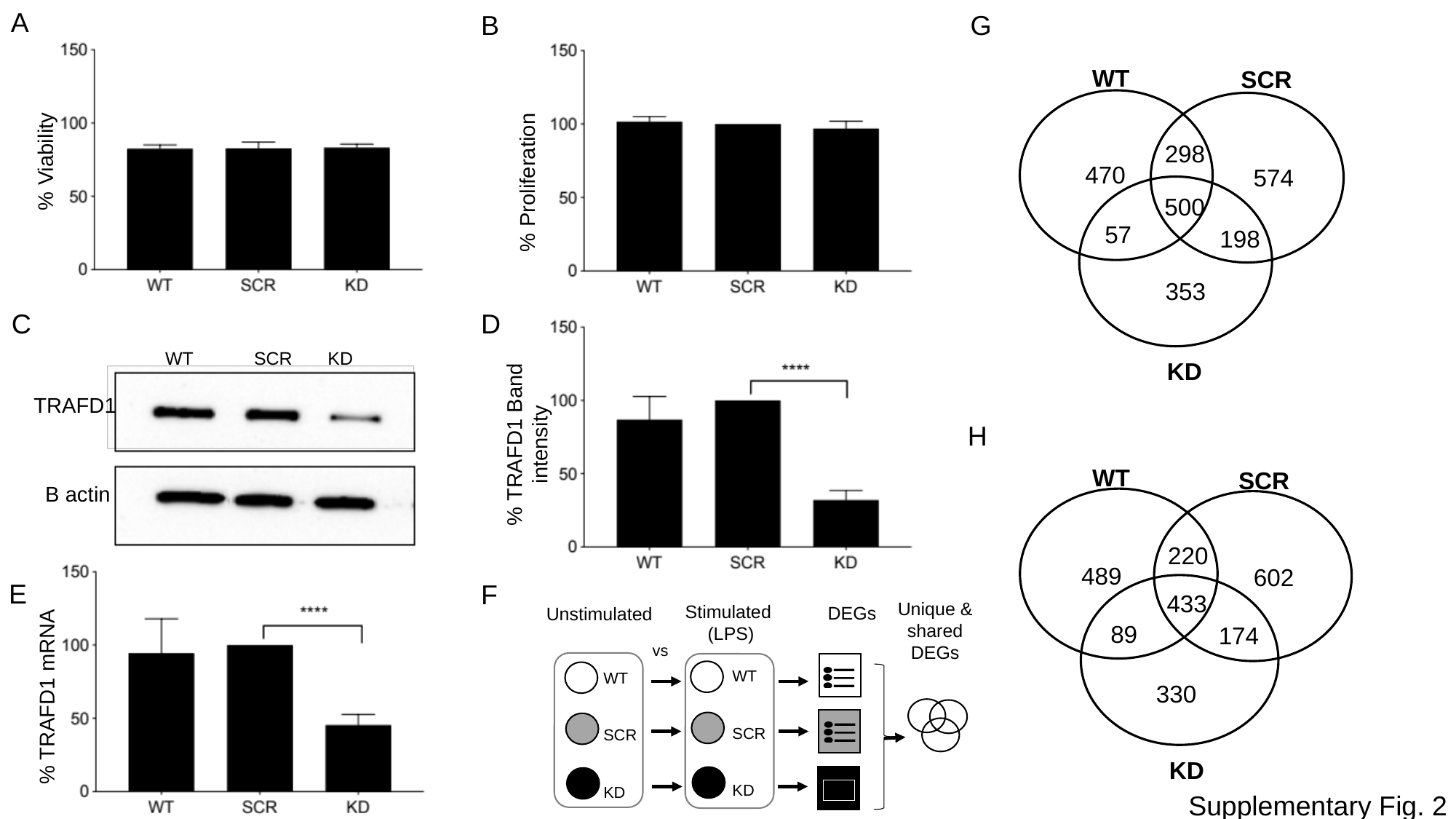

A
B
G
WT
SCR
298
470
574
500
57
198
353
KD
% Viability
% Proliferation
D
C
 WT SCR KD
TRAFD1
% TRAFD1 Band intensity
H
WT
SCR
220
489
602
433
89
174
330
KD
B actin
E
F
Unique & shared DEGs
Stimulated
(LPS)
DEGs
Unstimulated
vs
WT
SCR
KD
WT
SCR
KD
% TRAFD1 mRNA
Supplementary Fig. 2

### Slide 3
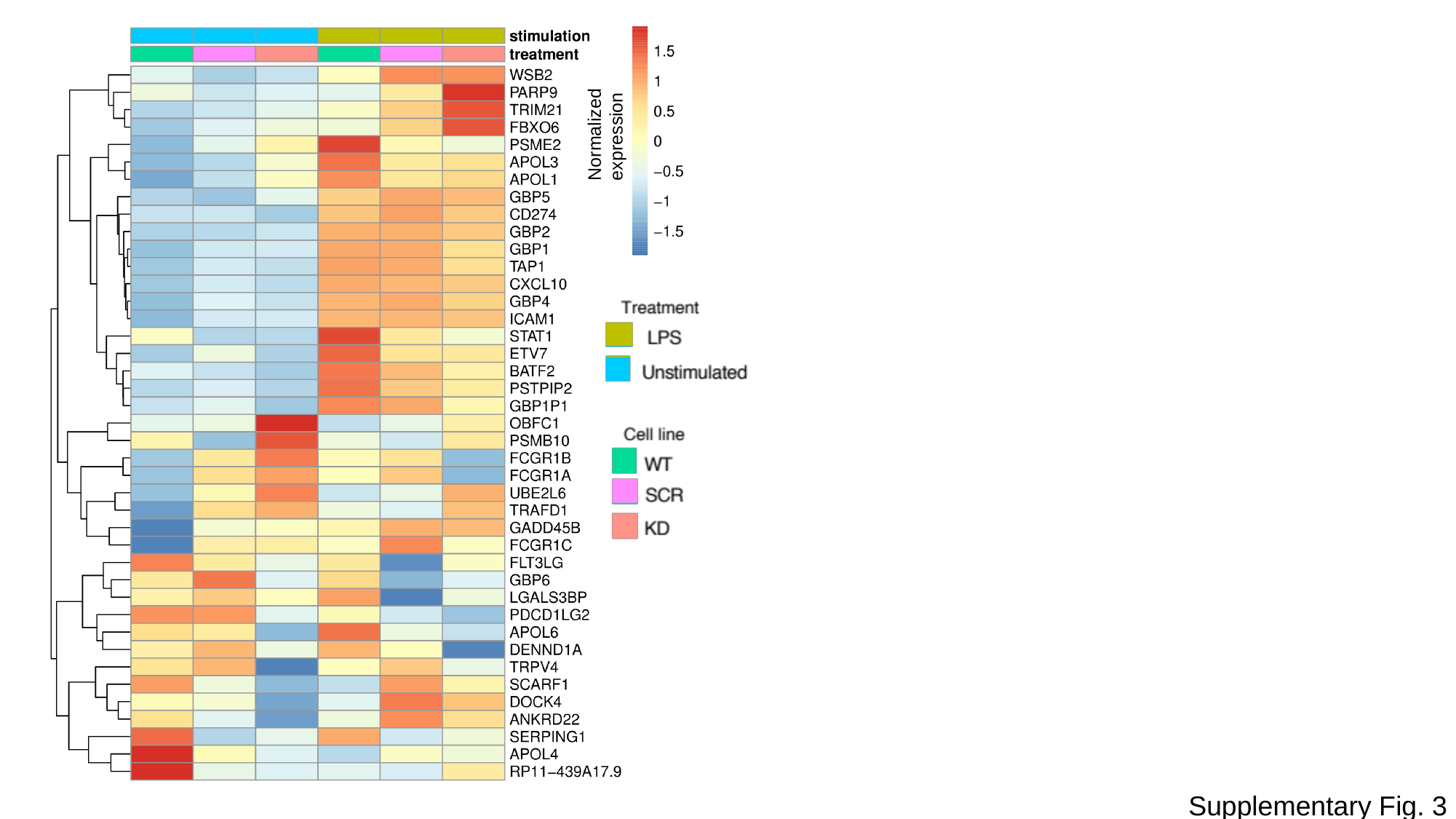

Normalized expression
Supplementary Fig. 3

### Slide 4
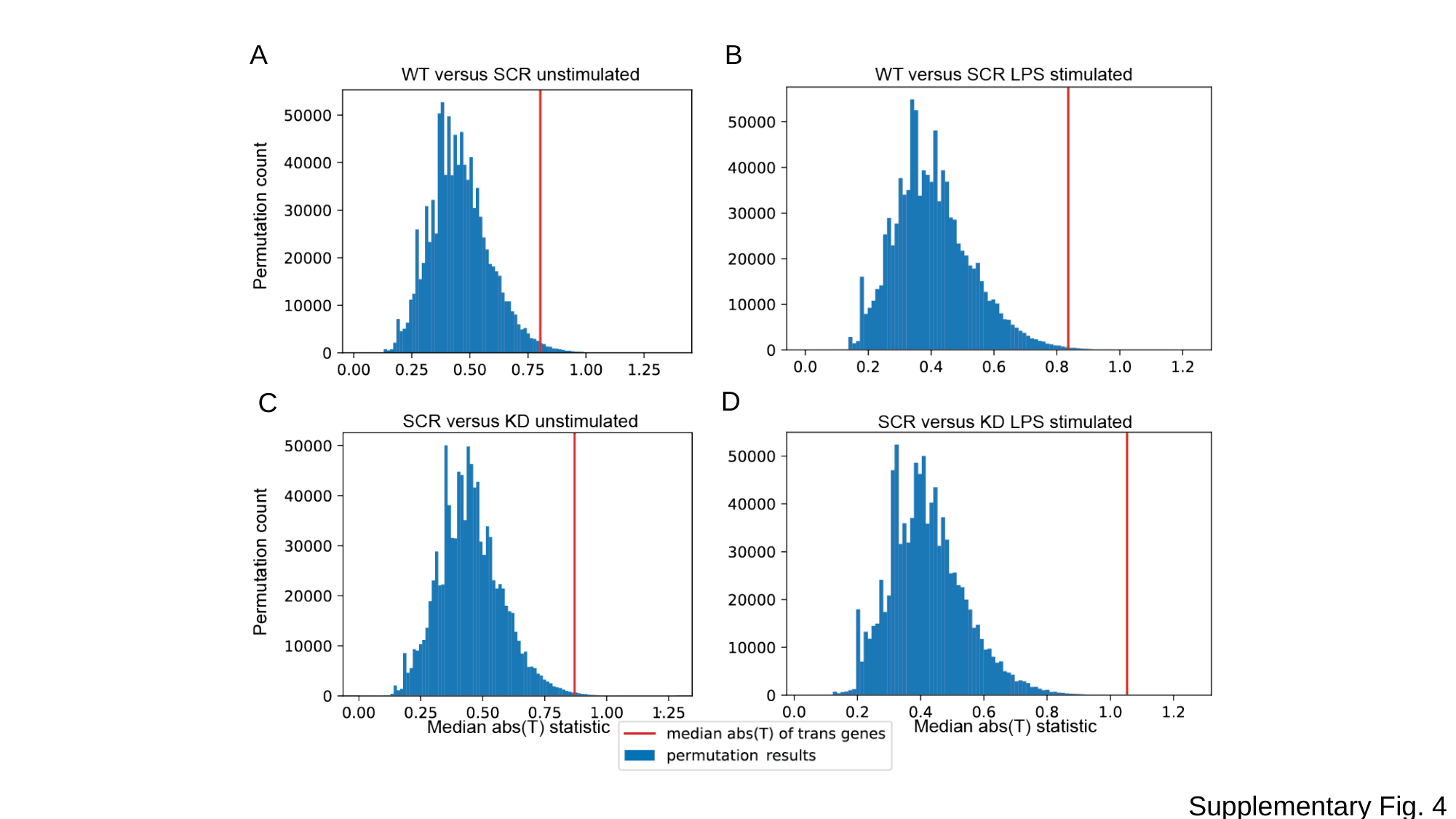

A
B
B
D
C
Supplementary Fig. 4
